# Phase separation of PGL-3 driven by structured domains that oligomerize and interact with terminal RGG motifs

**DOI:** 10.1101/2025.06.23.660947

**Authors:** Rimpei Kuroiwa, Piyoosh Sharma, Andrea Putnam, Stephen D. Fried, Geraldine Seydoux

## Abstract

Phase separation of biomolecular condensates is often assumed to be driven by interactions involving nucleic acids and intrinsically disordered regions (IDRs) of proteins. PGL-3 is a component of P granules, biomolecular condensates in the *C. elegans* germline, that contains two structured domains in tandem (D1-D2), an internal IDR, and a C-terminal IDR rich with RGG motifs. Theoretical and *in vitro* studies have implicated the internal IDR and RGG motifs in driving PGL-3 phase separation via self-interactions and binding to RNA. Studies in cells, however, have implicated the D1 and D2 domains. Here, we investigate the molecular basis of PGL-3 phase separation *in vitro* using microscopy, crosslinking mass spectrometry and biophysical measurements. We find that D1-D2 is oligomeric and necessary and sufficient for phase separation independent of RNA. D1-D2 also interacts with the terminal RGG domain in a manner that correlates with phase separation. In contrast, the internal IDR is neither necessary nor sufficient for phase separation. These findings support a new model for PGL-3 phase separation driven by oligomerization of structured domains and enhanced by RGG repeats independent of RNA.

## Introduction

Biomolecular condensates are proposed to assemble by liquid-liquid phase separation (LLPS), a thermodynamically driven de-mixing phenomenon ^1–3^. Several studies have highlighted the role of nucleic acids and intrinsically disordered regions (IDRs) of proteins in driving phase separation. Studies focused on FUS and TDP-43 revealed that low-complexity regions (LCRs), a subclass of IDRs, provide weak, multivalent interactions, including cation-π interactions and π-π stacking, that drive LLPS ^4^. These studies have motivated the development of predictors that estimate LLPS propensity based on primary sequence with no or limited three-dimensional structural information ^5–11^.

Phase separation of IDR-containing proteins can also involve folded domains^12^. For example, TDP-43 condensation is enhanced by the head-to-tail oligomerisation of its N-terminal domain ^13,14^. Folded domains can also provide binding surfaces for short linear motifs (SLiMs) in IDRs^15^, as in the case of Rubisco which phase separates with the disordered EPYC1 protein in pyrenoids ^16–18^, and NCK whose SH2 and SH3 domains interact with proline-rich motifs (PRMs) in N-WASP and pTyr residues in Nephrin, respectively ^19,20^. Synthetic reconstitutions have also shown that dimerization and oligomerisation induced by folded domains can drive LLPS by bringing together self-interacting multivalent IDRs^21–23^. In the case of G3BP, a N-terminal dimerization domain and a C-terminal RNA-binding domain collaborate to drive phase separation in the presence of RNA^24^.

Whether structured domains can drive phase separation in the absence of IDRs or nucleic acids is less clear. A molecular dynamics simulation study reported that two multimeric coiled-coil (CC) domains connected by a linker segment would be sufficient for phase separation ^25^. SPD-5, a *C. elegans* pericentriolar material (PCM) scaffold protein that contains nine CC domains separated by mostly disordered linkers, can form glass-like or gel-like condensates in the presence of a crowding agent *in vitro* ^26^. Two coiled-coil domains, a leucine zipper (LZ) domain, and Cnn-motif 2 (CM2) domains from *Drosophila* centrosomin (Cnn, a functional orthologue of SPD-5) were reported to come together in micron-scale assemblies when mixed *in vitro* ^27^. Similarly, a synthetic study showed that two CC domains fused to GFP by flexible linkers can form liquid-like condensates in cells and in the presence of a molecular crowder *in vitro* ^28^. These examples suggest that certain folded domains can drive condensation on their own.

PGL proteins were the first proteins identified as constitutive components of P granules, biomolecular condensates in the *C. elegans* germline^29,30^. Early studies reported that PGL-1 and its paralog PGL-3 are required for P granule assembly in embryos and form cytoplasmic granules when expressed individually or together in CHO cells^31,32^. Crystallography and size exclusion chromatography of residues 1-212 and residues 205-447 in PGL-1 revealed that each form well-folded homodimerization domains (D1 and D2), separated by a likely disordered six amino-acid linker ^32,33^. PGL-1 and PGL-3 are homologous with 60% identity overall and contain, in addition to the N-terminal D1 and D2 domains, an internal IDR (223 residues in PGL-1, and 175 residues in PGL-3) followed by a C-terminal, low sequence-complexity domain rich in arginine (R) and glycines (G), including 10 RGG and 1 RG motifs in PGL-1 and 6 RGG and 6 RG motifs in PGL-3 (Figs. 1A, SI Appendix, Fig. S1A-B). We refer to this C-terminal domain as the “RGG domain”.

**Figure 1.**
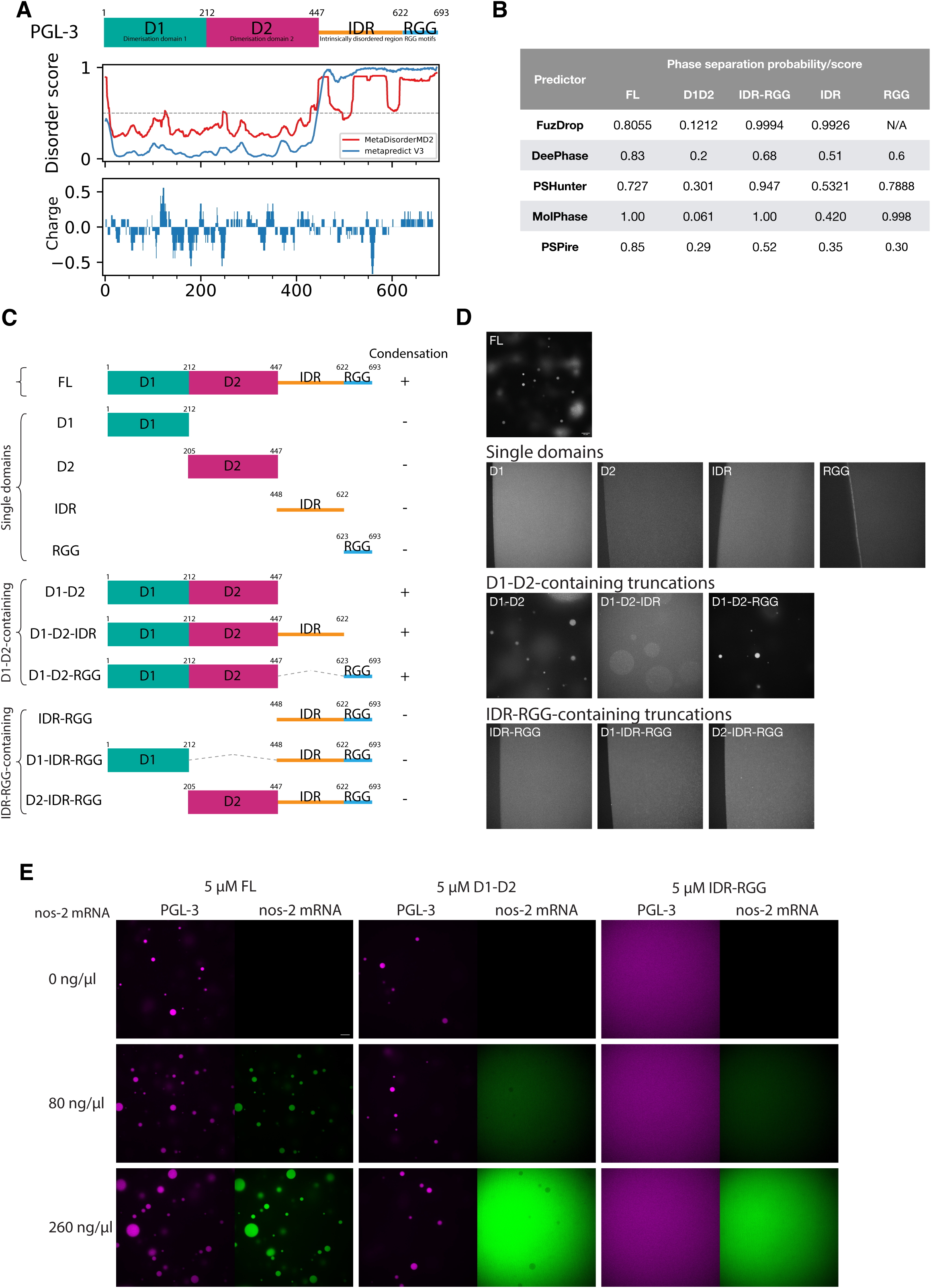
The N-terminal D1-D2 domain is necessary and sufficient for phase separation. (A) Schematic showing the domain architecture of PGL-3 aligned to disorder scores predicted from MetaDisorderMD2^61^ and metapredict V3^62^, and a charge plot by EMBOSS charge ^70^. (B) Table showing the phase separation probabilities predicted for full-length, the ordered region of PGL-3 (D1-D2) and the disordered region of PGL-3 (IDR-RGG), as well as IDR and RGG separately, using FuzDrop^5^, DeePhase^6^, PSPHunter^7^, MolPhase^8^ and PSPire^9^. The FuzDrop score for the RGG domain shows N/A as FuzDrop required a longer input sequence. (C) Schematics showing PGL-3 fragments tested for condensation *in vitro* as shown in D. (D) Fluorescence micrographs of condensation reaction mixtures. The indicated PGL-3 domains (20 μM, 1% trace labelled with Alexa Fluor 647) were mixed in 67.5 mM NaCl, 25 mM HEPES pH 7.5 on a glass bottom dish at 19°C for 10 min. In micrographs with no condensates, the air-water interface is shown on the left to highlight the homogeneous distribution of fluorescence in solution. Scale bar=10 μm. (E) Fluorescence micrographs of 5 µM PGL-3 and derivatives (<1% trace labelled with Alexa Fluor 647) with varying concentrations of *nos-2* mRNA (trace labelled with DyLight 488) in 125 mM NaCl. To highlight the distribution of labelled components, fluorescence intensities were adjusted for each protein and are comparable only across each column (micrographs containing the same protein with varying amounts of RNA). Scale bar=10 μm.

Different studies have pointed to different PGL domains driving condensation. A deletion spanning D1 and D2 prevented PGL-3 condensation in CHO cells and *C. elegans* embryos^31^, and point mutations in the D1 dimerization interface prevented PGL-1 condensation in CHO cells and in the *C. elegans* germline^32^. In addition, a preprint reported that the first 452 amino acids of PGL-3 (spanning D1 and D2) were sufficient for phase separation *in vitro* and that a C-terminal fragment (a.a. 370-693) failed to form condensates^34^. A theoretical study, however, found that a region spanning a portion of the internal IDR and the RGG domain (a.a. 515-693) was sufficient to reproduce the experimental phase diagram of full-length PGL-3 across varying temperatures and PGL-3 and salt concentrations^35^. Finally, a study using recombinant PGL-3::GFP showed that mRNA enhances PGL-3 condensation in a manner dependent on the RGG domain, which could bind RNA *in vitro* ^36^. Deletion of the RGG domain, however, did not prevent PGL-3 condensation in transfected CHO cells and *C. elegans* embryos^31^. In other systems, disordered RGG motif-containing regions have been reported to mediate binding to nucleic acids ^37–39^ and proteins, using homotypic RGG:RGG interactions ^40,41^ and heterotypic interactions involving other disordered domains ^24,42^ or folded domains ^43–45^.

To complement these studies, we have undertaken a systematic analysis of PGL-3 phase separation *in vitro* to determine how each domain contributes to the phase separation properties of PGL-3 in the absence of RNA. We find that PGL-3 phase separation is driven primarily by oligomerization of D1 and D2 domains and tuned by interactions between the RGG domain and D1-D2.

## Results

### Computational models predict high LLPS propensity for the IDR-RGG domain of PGL-3

To investigate which regions in PGL-3 promote phase separation, we first analysed full-length PGL-3 and its sub-domains, using five computational models trained to estimate LLPS propensity based on sequence (FuzDrop^5^, DeePhase^6^, PSPHunter^7^, MolPhase^8^, PSPire^9^). All use features derived from the primary sequence for prediction. PSPire in addition considers surface exposed residues in structured domains. All five models gave full-length PGL-3 a high propensity score for phase separation (0.7 or higher, where 0.5 is the threshold for binary prediction) (Fig.1B). Comparing sub-regions of PGL-3, the models scored the IDR-RGG domain highest and the D1-D2 domain lowest, consistent with the models relying primarily on protein disorder to predict phase separation.

### The D1-D2 domain is sufficient for phase separation

To experimentally determine the regions in PGL-3 responsible for phase separation, we turned to an *in vitro* reconstituted system. We purified from *E. coli* full-length, untagged PGL-3 and several derivatives and obtained a synthetic peptide spanning the RGG domain (Methods; Fig. 1C and SI Appendix, Fig. S1C-D). Each protein preparation was trace labelled (<1 %) with Alexa Fluor 647 to facilitate condensate imaging by fluorescence microscopy. Condensation was examined in solutions containing 20 µM protein, 67.5 mM NaCl, 25 mM HEPES pH 7.5 in the absence of RNA and crowding agents (Methods). Only full-length PGL-3 and derivatives that contained both the D1 and D2 domains yielded condensates (Fig.1C-D). Derivatives that contained only D1 or D2 failed to form condensates. Derivatives that contained the IDR and/or the RGG but lacked D1-D2 also failed to condense, even when tested at higher protein concentration (SI Appendix, Fig. S1E).

RNA enhances phase separation of full-length PGL-3 ^36^. As expected, when we supplemented the condensation reactions with RNA, we observed that RNA is enriched in condensates formed by full-length PGL-3 (Fig. 1E). RNA, however, did not enrich in D1-D2 condensates. The IDR-RGG domain did not form condensates in the presence of RNA (Fig.1E), even when tested at 100 µM (SI Appendix, Fig. S1F).

Aoki and co-workers ^32^ reported that mutations (R123E, K126E and K129E) that disrupt D1 dimerization block PGL-1 phase separation in tissue culture cells and in *C. elegans* germlines. We found that the same mutations introduced in full-length PGL-3 were sufficient to abrogate condensation at 10 µM and 30 µM protein and yielded only rare condensates when tested at 50 µM (SI Appendix, Fig. S1G). We conclude that the D1-D2 region of PGL-3 is necessary and sufficient for phase separation.

### D1-D2 exhibits a higher c_sat_ and lower condensate viscosity compared to full-length PGL-3

We compared the phase diagram of full-length PGL-3 and D1-D2 across protein and salt concentrations using solution turbidity (Fig. 2A, SI Appendix, Fig. S2A) and direct observation of condensates by microscopy (Fig. 2B, SI Appendix, Fig. S2B). Both assays revealed that, at a given salt concentration, condensation of D1-D2 requires higher protein concentrations than full-length PGL-3. In a phase separation regime, the protein concentration in the dilute phase (c_dil_) equals the saturation concentration (c_sat_) and is independent of input concentration. As expected, we found that D1-D2 exhibited a higher c_dil_ compared to full-length PGL-3 at the three salt concentrations tested (SI Appendix, Fig. S2C).

**Figure 2.**
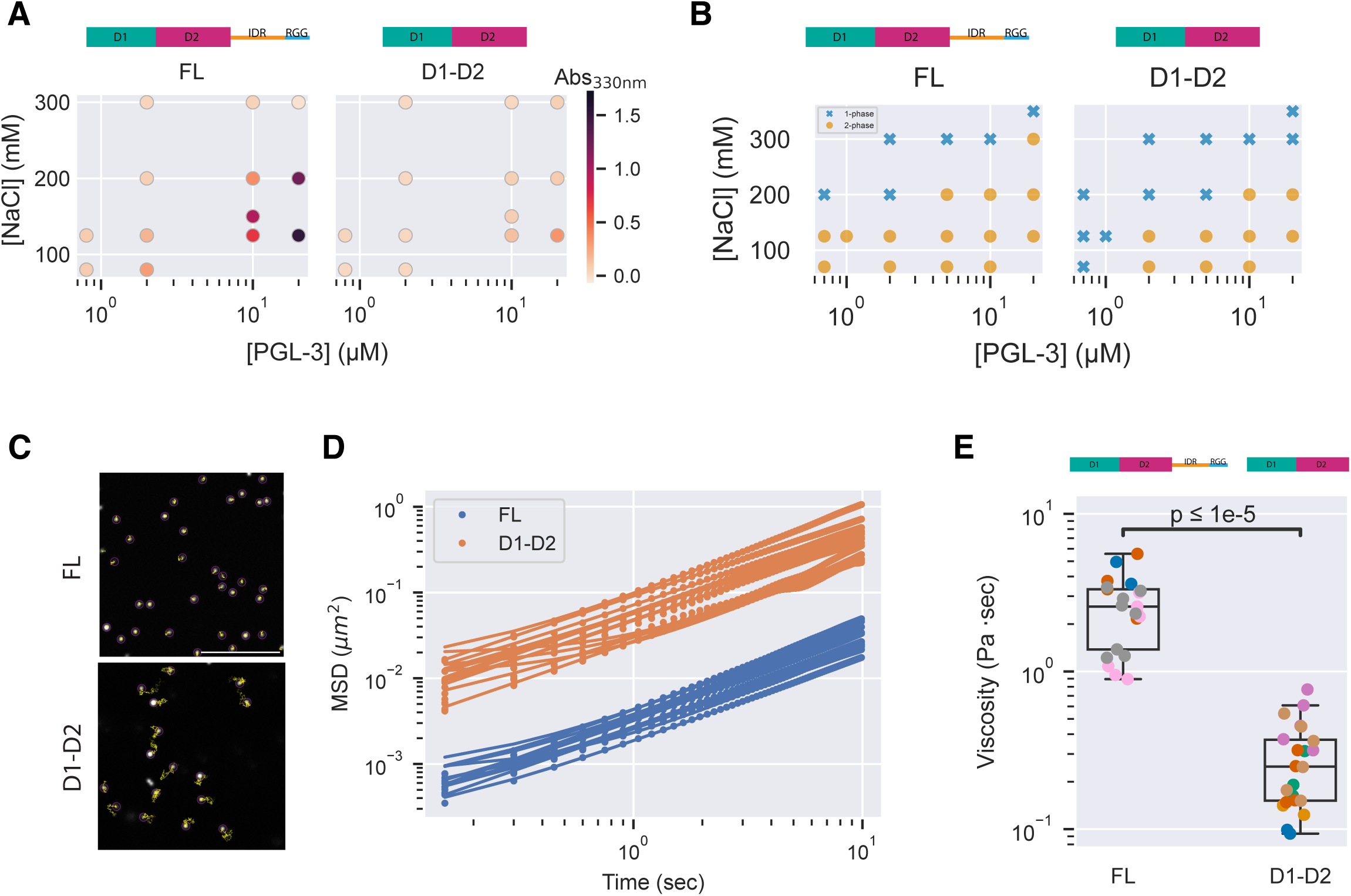
Properties of condensates assembled with full-length PGL-3 or D1-D2. (A) Plots showing the turbidity (absorbance at 330nm) of solutions containing full-length PGL-3 or D1-D2 at the indicated protein and salt concentrations in 25 mM HEPES pH7.5, <2.7 % glycerol at 19°C. Values shown are the average of three replicates. (B) Plots showing the presence (2-phase, dots) or absence (1-phase, crosses) of condensates visualized by fluorescent microscopy in solutions containing full-length PGL-3 or D1-D2 trace-labelled with Alexa Fluor 647 in 25 mM HEPES pH7.5, <2.7 % glycerol at 19°C. (C) Micrographs showing the trajectories over 15 seconds of polystyrene beads (0.2 μm) embedded inside a full-length PGL-3 or D1-D2 condensate. Each image shows a close-up of beads inside one condensate. Scale bar is 5 μm. (D) Plot showing the mean squared displacement (MSD) of beads diffusing inside full-length PGL-3 or D1-D2 condensates. Each line corresponds to the mean of MSDs from one condensate. (E) Plot showing the estimated viscosity based on measurements shown in C and D. Diffusion coefficient calculated from MSD were applied to the Stokes-Einstein-Sutherland equation to estimate the viscosity. Each dot represents a single condensate; colors designate condensates examined on the same day. P value calculated by Wilcoxon rank-sum test.

To estimate viscosity, we embedded fluorescent beads in the condensates and measured their diffusion by single particle tracking (SI Appendix, Fig. S2D and Methods). Beads in D1-D2 condensates exhibited larger mean square displacements (MSD) compared to beads in full-length PGL-3 condensates (Fig.2C, D, Movie S1). Using the Stokes–Einstein–Sutherland equation (D = k_B_T/6πηa), we estimated the viscosity of full-length PGL-3 condensates to be η = 3.30 ± 0.57 Pa·sec, consistent with values previous reported in the literature^30,46,47^, compared to 0.25 ± 0.09 Pa·sec for D1-D2 condensates (Fig.2E). We conclude that D1-D2 condensates are one order of magnitude less viscous than full-length PGL-3 condensates, suggesting that the IDR and/or RGG domain modulate the properties of PGL-3 condensates.

### The RGG domain enhances PGL-3 condensation

To examine possible contributions of the IDR and RGG domain to phase separation directly, we compared c_sat_ estimates for derivatives that contain D1-D2 with the IDR or RGG domain (Methods). At 125 mM NaCl and 5 µM PGL-3, D1-D2 formed droplets but D1-D2-IDR did not (Fig.3A). C_dil_ of D1-D2-IDR varied proportionally to input protein concentrations confirming lack of phase separation (Fig.3B; the apparently lower c_dil_ values are likely due to adsorption to tube surfaces). We conclude that the IDR domain reduces the phase separation propensity of D1-D2 when present in the absence of the RGG domain.

**Figure 3.**
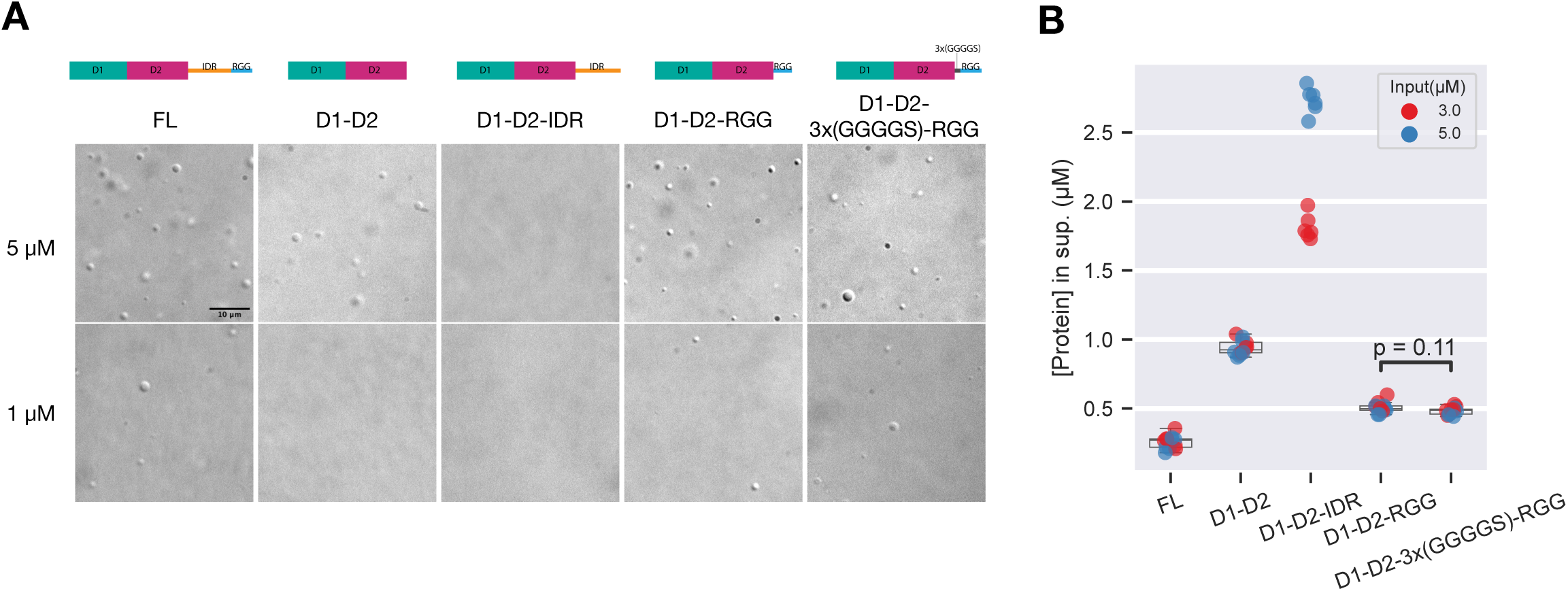
The RGG domain enhances D1-D2 condensation. (A) DIC micrographs of solutions containing FL and D1-D2-containing derivatives of PGL-3 at 1 or 5 µM, 125 mM NaCl. Scale bar=10 μm. (B) Plot showing the measured protein concentrations of the dilute phase with 3 µM (red) and 5 µM (blue) input PGL-3 concentrations in 125 mM NaCl. P-values calculated by Wilcoxon rank-sum test with Benjamini-Hochberg adjustment for multiple comparisons.

In contrast, we found that the RGG domain enhanced phase separation even in the absence of the IDR domain. D1-D2-RGG formed condensates at 1 and 5 µM PGL-3, 125 mM NaCl (Fig. 3A). The c_dil_ of D1-D2-RGG was 0.51 ±0.02 µM, lower than D1-D2 (0.94 ± 0.03 µM) (Fig.3B and SI Appendix, Fig. S3A) but higher than full-length PGL-3’s c_dil_ (0.26 ± 0.03 µM). Inserting a 3x(GGGGS) flexible linker between D1-D2 and the RGG domain did not significantly change c_dil_ (Fig.3A-B), possibly due to its shorter length (15 a.a.) compared to the IDR (175 a.a.). We conclude that the RGG domain enhances condensation of D1-D2.

### Crosslinking mass spectrometry reveals multiple interactions between the RGG domain and D1 and D2

To directly assess molecular interactions that accompany PGL-3 condensation, we performed crosslinking mass spectrometry (XL-MS) under salt regimes permissive and non-permissive for condensation (125 mM and 500 mM NaCl, respectively). We classified crosslinks within D1 and D2 into five categories based on their compatibility with SWISS-MODEL^48^ models of the PGL-3 D1 and D2 domains templated on the published dimer structures of isolated PGL-1 D1 and isolated PGL-1 D2 (SI Appendix, Fig. S4A) ^32,33^, as well as the monomer structures extracted from these. Crosslinks were considered “compatible” with monomer or dimer structures (intra-chain or inter-chain, respectively) if alpha carbons of the corresponding residues were less than 30Å apart in those models (SI Appendix, Methods).

We detected several homodimer crosslinks (Fig. 4A, yellow downward-pointing loops), which we define as crosslinks that occurred between the same residue on peptide-pairs that overlap in sequence. Because each tryptic PGL-3 fragment is unique, homodimer crosslinks must derive from two interacting PGL-3 molecules. D1:D1 homodimer crosslinks at K134 and D2:D2 homodimer crosslinks at K352 were compatible with the published D1 and D2 dimer structures ^32,33^ (Fig. 4A, yellow downwards loop crosslinks). Other D1:D1 and D2:D2 homodimer crosslinks were incompatible with published dimer structures: K5 and K277 (observed in both conditions), S272 (observed only under the condensing condition), and S434 (observed only under the non-condensing condition) (Fig.4A, SI Appendix, Fig. S4B, magenta crosslinks). These crosslinks suggest that the D1 and D2 domains contain at least one additional homotypic binding interface.

**Figure 4.**
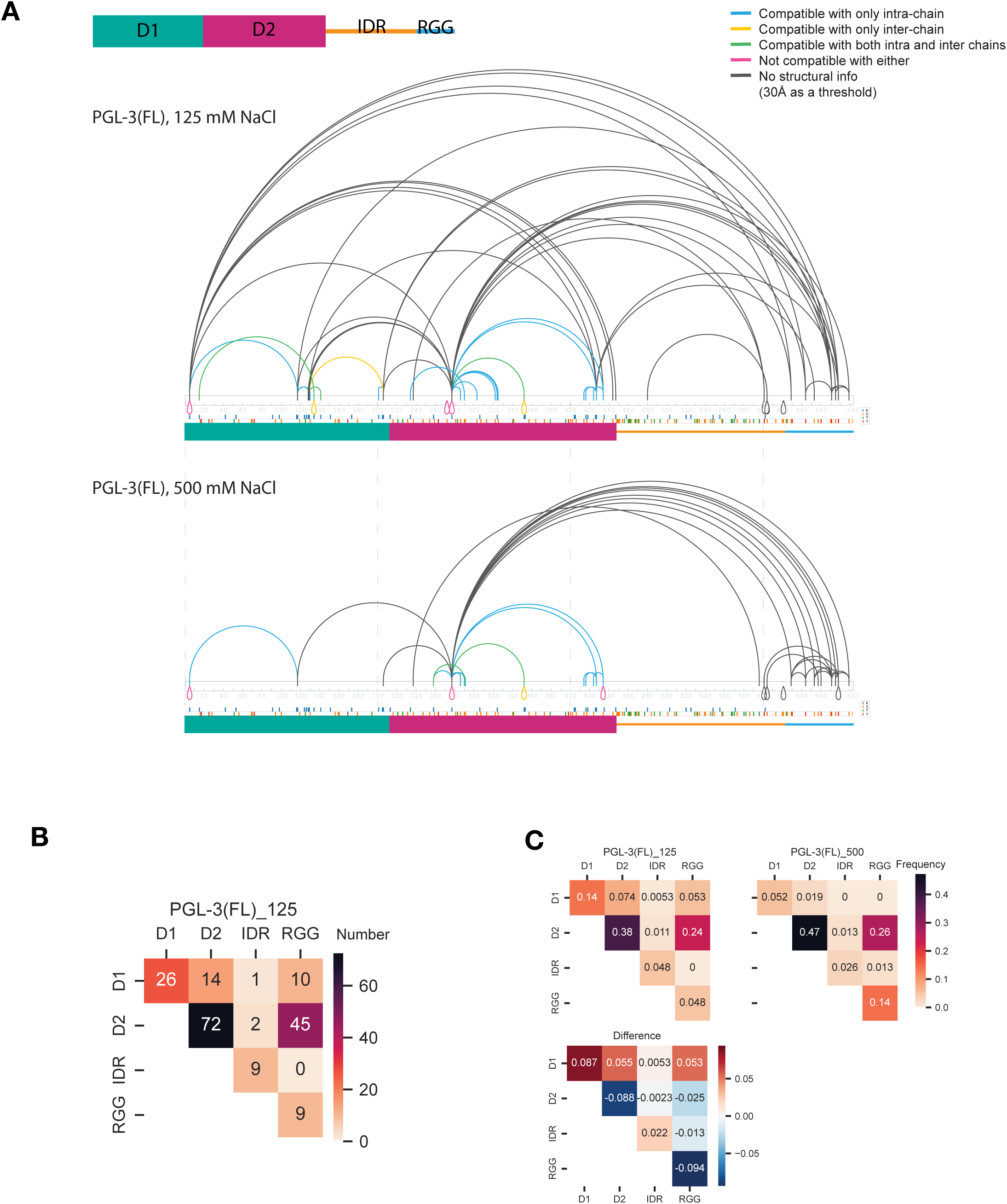
Intra- and inter-domain interactions revealed by XL-MS. (A) Connectograms showing all residue pairs identified by XL-MS on full-length PGL-3 in 125 mM NaCl (condensing condition) or 500 mM NaCl (non-condensing condition). Different colors indicate compatibility of each crosslink to structural models if the residue pair is intra-chain (monomer) and/or inter-chain (dimer) ^32,33^. Blue: compatible with intra-chain only, yellow: compatible with inter-chain only, green: compatible with both, red: not compatible with either, gray: no structural information. (B) Crosstab heatmap showing the number of crosslinked peptide-spectrum matches (XSMs) whose connecting residues reside within or between the four indicated domains at the 125 mM NaCl (condensing) condition. (C) Crosstab heatmaps showing the frequency of XSMs whose connecting residues reside within or between the four indicated domains. Top two heatmaps show cross links obtained at 125 mM (permissive for condensation) and 500 mM NaCl (non-permissive for condensation). The crosstab titled “difference” shows values calculated by substracting 500 mM NaCl values from 125 mM NaCl values; positive values (red colors) indicate crosslinks obtained at higher frequencies under condensing conditions.

We also detected many crosslinks between peptides in different domains of PGL-3 (“inter-domain” crosslinks) (Fig.4A). Unlike homodimer crosslinks, it is not possible to distinguish whether inter-domain crosslinks occurred intra- or inter-molecularly. We detected 18 spectra corresponding to crosslinks that connect residues within the IDR-RGG region and 58 spectra corresponding to crosslinks that connect residues in the D1-D2 region to residues in the IDR-RGG region. Among the latter, 55 involved the RGG domain and only 3 involved the IDR (Fig.4B). Remarkably, all the crosslinks between D1 and the RGG domain appeared only under condensing conditions (Fig. 4A). We systematically compared the frequency of each crosslink class under the two salt concentrations (Fig. 4C). This analysis revealed that D1:RGG and D1:D crosslinks are favored under condensing conditions. In contrast, D2:D2, D2:RGG, IDR:RGG and RGG:RGG crosslinks are favored under non-condensing conditions (Fig. 4C). Crosslinks involving the IDR are the rarest crosslink type.

Together the XL-MS findings suggest that the D1 and D2 domains are multivalent, possibly forming oligomers, and interact with the RGG domain. The internal IDR, in contrast, rarely participates in intermolecular interactions either with itself or with other PGL-3 domains.

### D1 and D2 form oligomers and interact with the RGG domain

To directly test whether D1-D2 forms oligomers, we examined dilute solutions (50 to 100 nM) of D1-D2 by mass photometry, a light-based technique that allows estimation of the mass of molecular complexes in solution (SI Appendix, Fig. S5A). We readily detected particles whose masses match the theoretical sizes of D1-D2 monomers, dimers, tetramers and hexamers, even at tens of nano-molar concentration range, well below the physiological concentration of PGL-3 (0.68 µM ^36^). As expected, the frequency of larger oligomers increased with concentration. These observations confirm that D1-D2 can form oligomers, consistent with work by Aoki and co-workers^33^, who observed D1-D2 trimers and/or tetramers using chemical crosslinking.

Individual D1 or D2 domains are too small by mass to resolve reliably by mass photometry. Therefore, to examine their oligomerization potential, we turned to chemical crosslinking using 1% formaldehyde (FA) and trace-labeled D1 and D2 (DL488). The crosslinked species were resolved by SDS-PAGE and examined by in-gel fluorescent imaging (Methods). We found that both D1 and D2 formed dimers and higher oligomeric complexes, confirming that these domains can form oligomers when present as single domains in solution (Fig. 5A).

**Figure 5.**
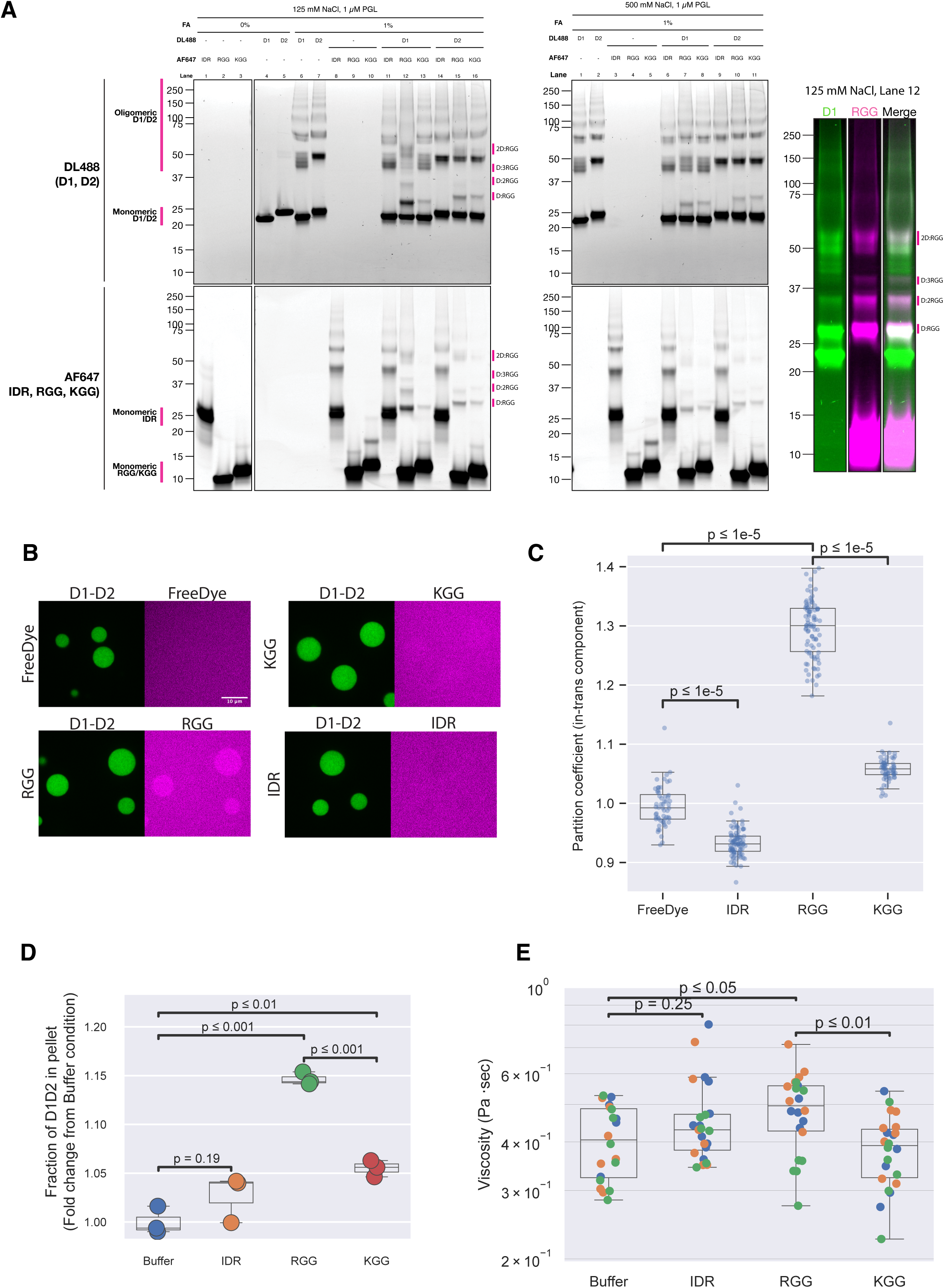
The RGG domain binds to D1 and D2 in *trans* and modulates D1-D2 phase separation. (A) Fluorescence images of SDS-PAGE gels showing crosslinked complexes of PGL-3 fragments. Top and bottom images were obtained from the same gel, scanned for the two different fluorophores. Top: D1 and D2 labelled with DyLight 488. Bottom: IDR, RGG and KGG labelled with Alexa Fluor 647. Color image is color version of lane 12 from the 125 mM NaCl condition. (B) Micrographs comparing partitioning of the IDR, RGG and KGG (magenta) to D1-D2 condensates (green). Results are quantified in C. (C) Plot showing the partition coefficients of component supplied in *trans* to D1-D2 condensates. Each dot represents a value from a single condensate. P-values calculated by Wilcoxon rank-sum test with Benjamini-Hochberg adjustment for multiple comparisons. (D) Plot showing the fraction of D1-D2 protein in pellet over total protein normalized to the buffer control, comparing different PGL-3 domains added *in trans*. P-values calculated by Wilcoxon rank-sum test with Benjamini-Hochberg adjustment for multiple comparisons. (E) Plot showing the estimated viscosity of D1-D2 condensates with the additional component indicated on the x-axis supplied in *trans*. Viscosity measurement was done by passive rheology (Methods). P-values calculated by Wilcoxon rank-sum test with Benjamini-Hochberg adjustment for multiple comparisons

To determine whether D1 or D2 also interact with the IDR or RGG domains, we trace labeled the IDR and RGG domains with a different fluorophore [Alexa Fluor 674 (AF647)] to detect possible hetero-oligomers. We also tested a KGG peptide where all the Arg residues in the RGG domain were replaced with Lys (SI Appendix, Fig. S5B). Mixing the IDR domain with D1 or D2 did not affect the migration patterns of either species, confirming minimal interactions between these domains (Fig. 5A). In contrast, mixing D1 with RGG led to a dramatic shift in the migration patterns of both D1 (labeled with DL488) and RGG (labeled with AF647). The novel bands were positive for both fluorophores, consistent with D1:RGG crosslinked species (Fig. 5A Right, SI Appendix, Fig. S5C). The apparent sizes of the crosslinked D1:RGG species were consistent with 1:1, 1:2, 1:3 and 2:1 stoichiometries, as well as higher order oligomers of unknown stoichiometries. Similarly, we also detected D2:RGG crosslinked species. Most of the heterotypic species were absent in 500 mM NaCl (Fig.5A), suggestive of electrostatic interactions. We also tested a scrambled RGG peptide in which all six RGG triplets were disrupted (SI Appendix, Fig. S5B; Methods). The scrambled RGG peptide crosslinked to D1 and D2 as efficiently as the native RGG peptide (SI Appendix, Fig. S5D), supporting the theory that the interaction is largely electrostatic. We conclude that that the RGG domain interacts with both the D1 and D2 domains, as was also observed in the XL-MS data.

### The RGG domain enriches in D1-D2 condensates when provided in *trans*

To examine whether D1-D2 condensates can recruit the RGG peptide in *trans*, we induced D1-D2 condensation in the presence of equimolar amounts of IDR, RGG and KGG peptides and determined their partition coefficients in D1-D2 condensates (Methods, SI Appendix, Fig. S5E). As a control, we used free fluorescent dye, which exhibited no enrichment or depletion into D1-D2 condensates, demonstrating minimal effects of the fluorophores on partitioning (Fig. 5B-C). We found that D1-D2 condensates could enrich the RGG peptide with a partition coefficient of 1.30±0.01. The scrambled RGG also enriched in the D1-D2 condensates (SI Appendix, Fig. S5F), in contrast to the KGG peptide, which was recruited less efficiently, and the IDR domain which was weakly excluded (Fig. 5B-C). We conclude that D1-D2 condensates can enrich the RGG peptide, consistent with binding between D1-D2 and the RGG domain in condensates.

### The RGG domain enhances phase separation and increases viscosity of D1-D2 condensates in *trans*

So far, our data demonstrate that the RGG domain enhances D1-D2 condensation in *cis* (Fig. 3), can be crosslinked to D1 and D2 domains (Figs. 4, 5A) and can be enriched in D1-D2 condensates when provided in trans (Fig. 5B-C). We wondered if heterotypic interactions between the RGG domain and D1-D2 are sufficient to modify condensate properties. To test this hypothesis, we first compared the proportion of D1-D2 in the dense phase in the presence or absence of the RGG peptide. We found that addition of the RGG peptide increased the fraction of D1-D2 in the pellet by ∼15 % compared to buffer control (Fig. 5D). We observed a similar effect using the scrambled RGG peptide and a weaker effect with the KGG peptide (Fig. 5A and SI Appendix, Fig. S5G). Addition of the IDR did not yield a statistically significant change (Fig.5D). We conclude that the RGG domain weakly enhances phase separation of D1-D2 when provided in trans.

Next, we tested whether the RGG domain also increases the viscosity of D1-D2 condensates. Using embedded beads, we measured the viscosity of D1-D2 condensates formed in the presence of equimolar IDR, RGG or KGG (Methods). Only the RGG increased condensate viscosity compared to buffer (by ∼1.25 fold, Fig. 5E). We conclude that interactions between the RGG and D1-D2 decrease c_sat_ and increase condensate viscosity.

### D1-D2 and RNA compete for binding to the RGG domain

The RGG domain of PGL-3 was previously shown to bind RNA in vitro^36^. To assess the relative interaction strengths of D1-D2:RGG and RNA:RGG interactions, we compared partitioning of the RGG peptide into D1-D2 condensates in the presence or absence of *in vitro* transcribed *nos-2* mRNA (SI Appendix, Fig. S5E). When tested at a physiological stoichiometry of PGL-3 and RNA calculated based on values reported previously^36^ (4 µM PGL-3 and 300 ng/µl *nos-2* RNA; Methods), the RGG domain still enriched in D1-D2 condensates although at a lower partition coefficient compared to the no-RNA condition (Fig. 6A-B). A higher RNA concentration of 500 ng/µl decreased partitioning of the RGG peptide into D1-D2 condensates, consistent with RNA competing with D1-D2 for RGG binding. The RGG peptide also formed aggregates with RNA in the presence or absence of D1-D2 (Fig. 6A and SI Appendix, Fig. S6A). We conclude that RNA and RGG competes for D1-D2 binding and that D1-D2:RGG interactions are maintained under physiological ratios of RNA, raising the possibility that these may be favored when the RGG domain is covalently linked to D1-D2 as in full length PGL-3.

**Figure 6.**
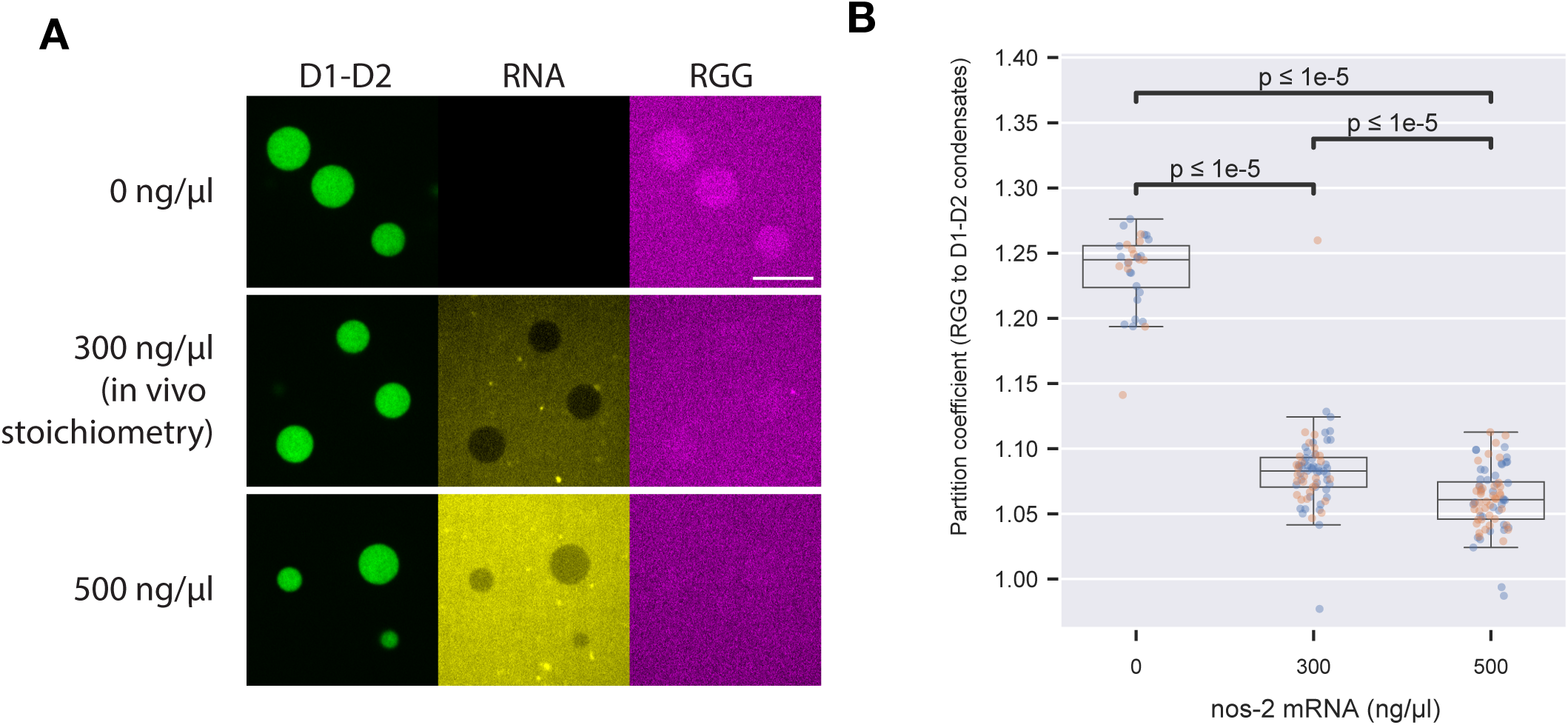
D1-D2 competes with RNA for binding to RGG domain (A) Fluorescence micrographs showing distribution of D1-D2 (green), *nos-2* mRNA (yellow) and the RGG peptide (magenta) upon D1-D2 condensation, at 4 µM D1-D2 and 4 µM RGG with varying concentrations of RNA. Quantification is show in B. (B) Plot showing the partition coefficients of the RGG peptide to D1-D2 condensates at varying concentrations of RNA. P-values calculated by Wilcoxon rank-sum test with Benjamini-Hochberg adjustment for multiple comparison.

## Discussion

In this study, we have characterized the molecular mechanism of phase separation by the P granule protein PGL-3. Contrary to computational predictions, we find that the folded domains of PGL-3 (D1 and D2) are necessary and sufficient for phase separation. Crosslinking experiments suggest that these domains confer multivalency to PGL-3, allowing it to form oligomers larger than dimers in solution and in condensates. D1-D2 also interact with the C-terminal RGG domain and these heterotypic (e.g. D1:RGG) interactions enhance phase separation and increase the viscosity of PGL-3 condensates. In contrast, the internal IDR was fully soluble under the conditions tested and is neither necessary nor sufficient for phase separation. Based on these observations, we propose the following working model for PGL-3 phase separation. D1 and D2 both have the ability to form oligomers and provide the requisite multivalency for condensation. The RGG domain further enhances phase separation by interacting in trans with D1-D2. The internal IDR domain contributes indirectly to phase separation by serving as a flexible linker between D1-D2 and RGG. We discuss aspects of this model in the following sections.

### PGL-3 condensation is driven primarily by oligomerization of the folded D1 and D2 domains

Our systematic screening of PGL-3 derivatives revealed that the first 447 amino acids of PGL-3, which span the folded domains D1 and D2, are necessary and sufficient for phase separation. Analyses of the D1 and D2 domains by X-ray crystallography revealed that both can form homodimers ^32,33^. Using XL-MS, we identified seven unique homodimer crosslinks in D1 and D2 compatible with the published homodimer structures ^32,33^, suggesting that the dimerization models observed in crystals are relevant in solution and in liquid condensates. We also identified four additional unique homodimer crosslinks that were not compatible with the published structures, suggesting that D1 and D2 also dimerize using interfaces not seen in the X-ray structures. Consistent with these findings, we detected homo-oligomers larger than dimers for both D1 and D2 using FA crosslinking in solution. We conclude that D1 and D2 are oligomerization domains that contain at least two homotypic binding interfaces that can be co-occupied on the same molecule. We suggest that D1 and D2 domains provide the multivalency required to create higher-order networks of PGL-3 molecules and are the primary drivers of PGL-3 phase separation.

### PGL-3 condensation is enhanced by interactions between D1-D2 and the RGG domain

Several lines of evidence indicate that D1-D2 also interact with the RGG domain. By XL-MS, we detected 10 and 45 spectra corresponding to crosslinks between the RGG domain and D1 and the RGG domain and D2, respectively, under phase separating conditions. The D1 domain, which is furthest away in sequence from the RGG domain, interacted with the RGG domain only under phase separating conditions in XL-MS experiments. We observed hetero-oligomers of various stoichiometries in crosslinked mixtures of RGG peptide and D1 or D2 domains. The RGG peptide is enriched in D1-D2 condensates, and lowers c_sat_ and increases condensate viscosity when attached covalently to D1-D2 (*cis* effect) as well as when added to D1-D2 condensates (*trans* effect, albeit more modestly).

We propose two related models for how the RGG domain tunes the phase separation properties of D1-D2. In the first model, binding between the RGG domain and D1-D2 simply increases total valency, which augments the PGL-3 network, lowers c_sat_ and increases condensate viscosity. In a second model, the RGG domain competes for one of the binding surfaces used by D1-D2 to oligomerize, which alters the landscape and kinetics of the network. In support of the second model, residues in D1 and D2 (K5 and K277) were detected by XL-MS in both homodimer crosslinks and heterotypic crosslinks involving the RGG domain (Fig.4A), raising the possibility that the same interface(s) that mediate D1 and D2 homodimer interactions also mediate D1-D2:RGG interactions. One possibility is that certain homotypic binding interfaces in D1-D2 lead to assemblies that are less capable of forming a dynamic higher-order network and therefore tend to terminate at soluble oligomers. Competition for these sites by the RGG domain could theoretically enhance condensation if oligomers involving the RGG domain were less soluble and/or more prone to forming high-order networks. A scrambled RGG peptide behaved similarly to the native RGG peptide in crosslinking to D1 and D2, enhancing the D1-D2 condensation and recruitment to D1-D2 condensates, suggesting that the specific sequence of the RGG domain is not essential, consistent with variability in the numbers of RGG motifs between PGL-1 and PGL-3. We also note that the effect of the RGG domain on condensation was considerably higher in *cis* than in *trans*. A likely possibility is that covalent linkage of the RGG to D1-D2 increases the effective k_on_ between RGG and D1-D2. We cannot exclude, however, the possibility that the RGG domain also enhances phase separation by forming RGG homo-oligomers as has been reported for other RGG domains^40,41^. Crosslinks within the RGG domain were recovered in the XL-MS data, but our experimental setup did not allow us to distinguish intra vs. inter molecule crosslinks. We attempted to generate point mutants that specifically blocked D1-D2:RGG interactions, but unfortunately these also interfered with phase separation of D1-D2 alone, potentially indicative of the dual use of binding interfaces as discussed above. The observation that the RGG domain impacts D1-D2 condensation *in trans* is strong evidence that the RGG domain does not function solely through RGG-RGG interactions.

Previous models of PGL-3 phase separation proposed that the RGG domain enhances PGL-3 phase separation by binding to RNA ^36^. Our competition assays indicate that RNA effectively competes with D1-D2 for binding to RGG, and that both binding interactions can be detected at stoichiometries of PGL-3 and RNA that mimic those *in vivo*. Whether the RGG domain of PGL-3 engages with RNA or D1-D2 *in vivo* is not known. Compared to other RNA binding proteins present in *C. elegans* embryos (e.g. MEX-5 and MEG-3), PGL-3 has a lower binding affinity for RNA^36,49,50^. Genetic analyses have demonstrated that mRNA recruitment to P granules in embryos depends on MEG-3 and does not require PGL-1 or PGL-3. iCLIP experiments comparing PGL-1 and MEG-3 immunoprecipitated from embryo lysates identified only 18 transcripts bound to PGL-1 compared to 657 transcripts bound to MEG-3 ^51^. These observations raise the possibility that PGL:RNA binding is limited *in vivo*, leaving the RGG motifs available for binding to D1-D2.

### The internal IDR of PGL-3 contributes to phase separation indirectly by acting as a linker between the D1-D2 and RGG domains

Our experimental findings indicate that the central IDR in PGL-3 is neither necessary nor sufficient for phase separation. XL-MS and FA crosslinking revealed few interactions involving the internal IDR domain, and D1-D2-IDR exhibited higher c_sat_ than D1-D2, suggesting that the internal IDR of PGL-3 is solubilizing. Consistent with this view, FINCHES ^52^ predicts that the internal IDR domain is mostly repulsive to itself and contains only small regions of weakly attractive patches (SI Appendix Fig. S7A). Deletion of the IDR domain in PGL-3 led to a minor increase in c_sat_ which could not be rescued by insertion of a short 3x(GGGGS) linker, raising the possibility that the linker function of the IDR is length-dependent. Consistent with this possibility, the IDRs of PGL-1 and PGL-3 are both relatively long (223 and 175 amino acids) despite low overall sequence identity (43%) compared to D1-D2 (69%) and the RGG domain (61%) (SI Appendix Fig. S1A-B). The IDR could also provide sites for protein modifications that could tune phase separation. Phosphorylation of PGL-1 and PGL-3 by the mTOR homologue LET-363 was reported to enhance phase separation *in vitro* ^53^. The same study showed that three out of five phosphorylation sites reside in the PGL-1 IDR domain. In PGL-3, phosphoproteomics studies identified eight phosphorylation sites all located in the IDR ^54–57^ . It will be interesting to explore the impact of these post-translational modifications on phase separation.

### Limitations of this study

Our study explored the molecular basis of PGL-3 phase separation using *in vitro* assays. Although our findings are consistent with observations that demonstrated the importance of D1-D2 for PGL condensation in cells, we did not investigate directly whether the interactions we uncovered *in vitro* also occur *in vivo*. A major challenge is that, unlike our assays which examined PGL-3 condensation in isolation, PGL-3 condensation *in vivo* occurs in the presence of dozens of other P granule proteins, many of which also contribute to P granule assembly^58^. In fact, a preprint reported that introducing other P granule proteins can modify the dynamics of PGL-3 co-condensates assembled *in vitro* ^34^. Another challenge is that our buffer conditions do not accurately mimic the properties and crowding of the *C. elegans* cytoplasm. Although we could observe D1-D2 oligomers at relatively low concentrations (50-100 nM), D1-D2 condensation required higher concentrations (∼1 µM), near the range estimated for PGL-1/3 concentration in embryos (1.3 µM^36^ for PGL-1 and 0.68 ^36^/0.98 µM^59^ for PGL-3). Finally, as mentioned above, PGL proteins are modified *in vivo* and these modifications could alter their phase separation properties ^53,60^. The reductionist approaches we used in this study uncovered a role for folded domains and RGG motifs in driving phase separation and provide hypotheses to explore further in the *in vivo* context.

## Materials and Methods

### Computational predictions and calculation of physicochemical parameters

Disorder predictions were performed on the MetaDisorder server^61^ and metapredict V3^62^, and the charge distribution was calculated on EMBOSS-charge^63^ with the window length= 9. Phase separation prediction was performed on their respective servers for DeePhase^6^, FuzDrop^5^, PSHunter^7^ and MolPhase^8^. For PSPire, the prediction was run locally following authors’ instructions^9^. Molecular weights, pIs and charges were calculated on Prot pi (https://www.protpi.ch).

### Protein preparation

The RGG peptide and its derivatives were synthesized chemically. Other PGL-3 derivatives were expressed as fusions with a N-terminal His-MBP purification tag in *E. coli* Rosseta2 (DE3) cells (Sigma). The fusions were purified and the His-MBP tag was removed through four purification steps (amylose affinity, reverse nickel affinity, heparin-affinity or anion exchange, size exclusion) following published protocols with modifications^64^ on an ÄKTA pure FPLC or using loose resins. For fluorophore labelling, amine-reactive Alexa Fluor 647 NHS ester (Invitrogen) and DyLight 488-NHS-ester(Thermo Fisher) were used. See SI Appendix for details.

### Condensation assay and phase diagrams

To prepare imaging sides, coverslips were sonicated in 50 % Isopropyl alcohol, blow-dried and stored avoiding dust at room temperature. Coverslips were mounted on Attofluor Cell Chamber (Invitrogen) or sealed onto LabTek Chamber Slide (Thermo Fisher), and 3 mm diameter x 1 mm depth CultureWell silicone gasket (Grace Bio-labs) was applied to create imaging wells. Buffers, proteins and RNA were mixed directly in an imaging well to induced condensation at the desired protein and NaCl concentrations. When indicated, fluorophore-labelled proteins were mixed to aid imaging, but only up to 1 % of the protein population were labelled to minimise artefacts. To create phase diagrams, fluorophore-labelled PGL-3 solutions were scored by microscopy for visible droplets five minutes after initial mixing.

### Turbidity measurements

Condensation of unlabelled PGL-3 was induced by dilution to varying concentrations of PGL-3 and NaCl in 25 mM HEPES pH 7.5. The solution was quickly mixed by pipetting up and down before mounting 2 µl of NanoDrop One (Thermo Fisher) spectrometer. Turbidity was measured via absorbance at 330 nm (Fig.S2A).

### C_sat_ estimation by BCA assay

Unlabelled PGL-3 were diluted to induce condensation in 20 µl reactions. Following the incubation at the room temperature (18-19°C) for 1 h, the solution was centrifuged at 21,000 g for 20 min at 19°C using a temperature-controlled centrifuge(Fig.S2C, Left). 18 µl from the supernatant was directly used to determine the protein concentration using Pierce™ BCA Protein Assay Kits (Thermo Fisher) according to the manufacture’s description. The reported values are mean ± 95% CI.

### Viscosity measurements

Viscosity was estimated by recording the diffusion of 0.2 µm diameter FluoSphere Carboxylate-Modified Microspheres (Invitrogen). Detailed experimental and analysis protocols are provided in SI Appendix.

### Crosslinking mass spectrometry (XL-MS)

Crosslinking reaction of PGL-3(FL) was performed by combining 7.5 µM protein in 25 mM HEPES pH 7.5, 125 or 500 mM NaCl with disuccinimidyl dibutyric urea (DSBU, ThermoFisher) crosslinker to 2 mM final concentration. DSBU crosslinks K, S, T and Y residues to each other. After quenching, proteins were digested, and the resulting peptides were desalted and reconstituted in 0.1% formic acid for separation and analysis on a UltiMate3000 UHPLC system (Thermo Fisher) coupled with a Q-Exactive HF-X Orbitrap mass spectrometer (Thermo Fisher). Spectra were searched in Scout ^65^. Detailed experimental and analysis protocols are provided in SI Appendix.

### FA crosslinking assay

D1 and D2 were trace-labelled with DyLight 488 and imaged in the Cy2 channel. IDR, RGG and KGG were trace-labelled with Alexa Fluor 647 and imaged in the Cy5 channel. PGL-3 fragments were diluted to 2 µM in 125 mM NaCl, 25 mM HEPES pH 7.5. For protein mixtures, each protein was diluted at 2 µM. Diluted protein solutions were incubated at room temperature for 10 min.

Formaldehyde(FA) (Fisher scientific) was diluted to 2 % in 125 mM NaCl, 25 mM HEPES pH 7.5. Equal volumes of 2 % FA were added to 2 µM PGL-3 solutions and mixed by pipetting up and down to reach 1 µM PGL-3 domains and 1 % FA. Following the incubation at room temperature for 20 min, the reaction was quenched by f.c. 2.5 mM Tris, 20 mM glycine for 5 min. Home-made SDS sample buffer and final 5 % BME(Sigma) was added and boiled for 7 min at 70°C and SDS-PAGE was performed. We used home-made SDS sample buffer without dyes to increase sensitivity. The composition of SDS sample buffer at 1x was as follows: 106 mM Tris HCl, 141 mM Tris base, 2 % SDS, 0.51 mM EDTA, 2.5 % glycerol. After the electrophoresis, gels were quickly rinsed in distilled water and imaged on Amersham Typhoon (Cytiva) for Cy2 and Cy5 channels.

### Pelleting assay

#### Tube preparation

To minimise protein absorbing to polypropylene tubes, 5 mg/ml BSA were placed in tubes and incubated at room temperature overnight. Before use, the tube was rinsed three times with reaction buffer [125 mM NaCl, 25 mM HEPES pH 7.5] to remove residual BSA.

#### Experiment

In-trans component (unlabelled IDR, RGG, KGG) and Alexa Fluor 647 trace-labeled PGL-3(D1-D2) were diluted to 6 µM each in 125 mM NaCl, 25 mM HEPES pH 7.5 and incubated at ambient temperature 18-19°C for 20 min. The protein mixture was centrifuged at 21,300 g for 20 min at 19°C using temperature-controlled centrifuge. The supernatant was collected by pipetting, and the pellet was resuspended in 1xLDS sample buffer (Thermo Fisher). The pellet and supernatant fractions were run on SDS-PAGE. Gels were quickly rinsed in distilled water and imaged on Amersham Typhoon (Cytiva) for the Cy5 channel. Analysis was performed on Fiji^66^ and the PGL-3 D1-D2 band intensities were measured using a published protocol^67^, and the fraction of D1-D2 in pellet was determined as pellet band intensity divided by the sum of pellet band and supernatant band intensities.

### Partition coefficient determination

#### Experiment

We used D1-D2 labelled with DyLight 488 and RGG, IDR, KGG, scrambled RGG labelled with Alexa Fluor 647, and nos-2 labelled with Alexa Fluor 546. For free dye, Alexa Fluor 647 NHS ester was quenched with glycine. Coversides were coated with PEG-silane as described here^68^ to prevent wetting and aid image analysis. For Fig.5B, C, 5 µM of each protein was mixed. For Fig.6A, B, 4 µM of each protein at indicated RNA concentration was used. For calculating in-vivo stoichiometric concentration of RNA, we used values reported here ^36^: 0.68 µM PGL-3 and 50 ng/µl mRNA assuming average length of 1.5 kb. This converts to 294.1 ng/µl RNA for 4 µM PGL-3. The nos-2 mRNA is 1.2 kb in length.

#### Analysis

Custom scripts were used for analysis. We trained a classifier for semantic segmentation using sklearn library. The classifier was trained using manually labelled masks, into three classes ‘Condensates’, ‘Out-of-focus condensates’ and ‘Background’, in the D1-D2 channel using select images from the analysed dataset. After running segmentation on the D1-D2 channel, ‘condensate’ class was used to define dense phase of each condensate. For each condensate, the periphery of the ‘condensate’ class was defined as ‘dilute phase’ to minimise the effect of uneven illumination. Finally, partition coefficients were calculated as a ratio between median of two regions, median of dense phase/ median of dilute phase in the fluorescence channel of interest (Fig.S5E)

Additional details and methods are provided in SI Appendix.

## Data Availability

The mass spectrometry proteomics data have been deposited to the ProteomeXchange Consortium via the PRIDE ^69^ partner repository with the dataset identifier PXD064592. List of all the residue pairs are available in SI Appendix Dataset S1. All other data needed to evaluate the conclusions in this paper are present in the paper and/or the SI Appendix.

## Acknowledgments

We thank Dr. Philip Mortimer at the JHU Mass Spectrometry Core facility and Dr. Mario A. Bianchet at the JHU Macromolecular Biophysics Core for technical assistance. R.K. is funded by the XDBio program at Johns Hopkins University and of the Funai Foundation for Information Technology. P.S. acknowledges support from the Albstein Foundation for brain research. S.D.F. acknowledges support from the NIH Director’s New Innovator Award (DP2-GM140926), from the National Science Foundation (MCB-2045844), from a Camille Dreyfus Teacher-Scholar Award, and from a Sloan Fellowship. G. S. is an investigator of the Howard Hughes Medical Institute.

