## Supplementary material for "Phase separation of PGL-3 driven by structured domains that oligomerize and interact with terminal RGG motifs": SI Appendix

**Supporting Information Text**

**Supplementary Methods**

**Molecular cloning**
Plasmids used in this study are listed in Table S1. Plasmids were prepared using standard molecular cloning procedures in *E. coli* DH5α (Sigma) using the pMAL-c2X vector backbone. PGL-3 sequences were fused at their N-termini with either a 6xHis-MBP-6xHis-TEV or a MBP-6xHis-TEV purification tag, with or without a linker between the TEV recognition sequence and PGL-3.

**Protein expression and purification**
*Protein expression:*
Transformed *E. coli* Rosseta2 (DE3) cells were grown in lysogeny broth (LB) overnight, the subcultured in terrific broth supplemented with 75 μg /ml ampicillin at 37°C, 250 rpm until reaching OD600 of ~1.0. The culture was cooled down at 16°C for 10 min and 1 mM isopropyl β-D-1-thiogalactopyranoside(IPTG) was added to induce protein expression overnight at 16°C with 250 rpm shaking. Cells were collected by centrifugation and the pellet were either directly used for purification or frozen and kept at -80°C until further use. Bacterial pellets were sonicated in lysis buffer [10 μg/ml RNaseA(QIAGEN), 1 tablet/10 ml cOmplete™ Mini(EDTA-free) (Roche), 250 mM NaCl, 0.4 M L-arginine, 10% (v/v) glycerol, 1 mM DTT, 12.5 mM HEPES pH 7.5].

*Amylose affinity chromatography:*
Lysed cells were cleared by centrifugation and filtration, and the lysate was bound to loose amylose resin (NEB) in a gravity column or MBPTrap HP 5 ml column (Cytiva). The resin was washed with MBPNaCl buffer [20% (v/v) glycerol, 500 mM NaCl, 50 mM HEPES pH 7.5, 1 mM DTT] and eluted with MBPNaCl buffer supplemented with 20 mM maltose.

*TEV cleavage and reverse nickel affinity chromatography:*
In-house purified 6xHis-TEV protease was added to protein elution fractions to cleave the tag at 16°C overnight. For home-made TEV protease, pET29b-10xHis-Super TEV was acquired from Addgene, the 10xHis tag was reduced to 6xHis for convenience, and purified following the published protocol[1]. The digested purification tag and undigested fractions were removed by reverse His-tag affinity chromatography on HisTrap HP 5 ml column (Cytiva).

*Heparin affinity or anion exchange chromatography:*
PGL-3 fragments were purified further using a HiTrap Heparin HP 5 ml column (Cytiva) (for constructs containing the RGG domain) or a HiTrap Q HP 5 ml column (Cytiva) (for constructs lacking the RGG domain). The flow-through from reverse nickel affinity chromatography was collected and diluted 5-fold to reach 100 mM NaCl with dilution buffer [40 %(v/v) glycerol, 0.1 % NP-40, 25 mM HEPES pH 7.5], loaded onto a column, washed with 20 %(v/v) glycerol, 100 mM NaCl, 25 mM HEPES pH 7.5, and eluted by gradient elution from 150 to 1000 mM NaCl.

*Size exclusion chromatography and storage:*
Protein fractions were run on HiPrep Sephacryl S-200 (Cytiva) in native buffer [20 %(v/v) glycerol, 500 mM NaCl, 25 mM HEPES pH 7.5, 0.5 mM TCEP] and dialyzed against native buffer. Proteins were concentrated using appropriately-sized Ultra Centrifugal Filters (Amicon), filtered with a 0.2 μm PES filter (MDI), aliquoted, flash frozen in liquid nitrogen, and stored at -80 °C. Protein

concentration of the final stock was determined by absorbance at 280 nm and protein purity was assessed by SDS-PAGE followed by Coomassie G250 staining (Fig. S1C).

**Preparation of chemically synthesized peptides**

RGG, KGG and scrambled RGG peptides were custom-synthesized by GenScript. In the KGG peptide all Arginine residues are replaced with Lys (Fig. S5B). The scrambled RGG peptide was designed to scramble the sequence of the native RGG peptide and disrupt all RGG triplets. However, it contains six RG doublets at non-native positions whereas the native RGG peptide contains six RGG triplets and six RG doublets (Fig. S5B). We used this scrambled RGG peptide that contains RG doublets after repeated failures to synthesize peptides disrupting all RGG and RG motifs. The lyophilized peptides were suspended and dialyzed against native buffer to remove residual salt from synthesis. Peptides were filtered using 0.2 µm PES filter (MDI) and flash frozen in liquid nitrogen and stored at -80°C. Protein concentration of the final stock was determined by absorbance at 280 nm.

**Protein labelling with fluorophore**

Proteins were diluted to 25 µM and mixed with ~50 µM Alexa Fluor 647 NHS ester(Invitrogen) or DyLight 488-NHS-ester(Thermo Fisher) in native buffer, and reacted for 1-4 h in the dark, with the exception of the KGG peptide, which was reacted in 20 %(v/v) glycerol, 500 mM NaCl, 50 mM phosphate buffer pH 6.5 at 4°C overnight to preferentially label the N-terminal amine group. Free dyes were removed by running the solution in Zeba™ Spin Desalting Columns, 7K MWCO (Thermo Fisher) three times. Labelled proteins were filtered using 0.2 µm PES filter (MDI), aliquoted, flash frozen in liquid nitrogen, and stored at -80 °C. Protein and dye concentrations were measured using Nanodrop One. The dye to protein ratio never exceeded 1.0, indicating less than one fluorophore per molecule on average. Note that RGG and KGG peptides were run through the desalting columns despite their smaller molecular weight; although the recovery was lower, the final yield was sufficient for our purposes.

**RNA synthesis**

*nos-2* mRNA was synthesised using mMESSAGE mMACHINE™ T7 Transcription Kit (Invitrogen) following the manufacturer's recommendation as described [2]. DNA template was PCR amplified from a plasmid containing *nos-2* cDNA sequence. ChromaTide™ Alexa Fluor™ 488-5-UTP or 546–14-UTP (Invitrogen) was included at 1/20 of the reaction volume to trace-label fluorescently. LiCl precipitated RNAs were dried, resuspended in nuclease-free water and the integrity was verified by denaturing agarose gel electrophoresis.

**Microscopy**

Microscopy was carried out at the ambient temperature of ~19 °C using a AXIO Observer (Zeiss) with a CSU-W1 SoRA spinning disk confocal system (Yokogawa) and an iXon Life 888 EMCCD camera (Andor) using SlideBook software (Intelligent Imaging Innovations). The SoRaD1 filter set was used for all imaging including fluorescent and DIC. Unless otherwise noted, images were taken using a 100 x objective lens at 100 ms exposure. For viscosity measurements, the time lapse images were acquired using 100 x objective with a 2.8 x relay lens at 3 ms exposure and 150 ms interval.

**Viscosity measurements**

*Experiment:*

Viscosity was estimated by recording the diffusion of 0.2 µm diameter FluoSphere Carboxylate-Modified Microspheres (Invitrogen) in PGL-3 condensates as described in [3] (Fig. S2D). PGL-3 condensation was induced in a buffer solution containing microspheres in 125 mM NaCl, 25 mM HEPES pH 7.5. Condensates were allowed to settle onto the coverslip surface for 5 min and all the image acquisition was finished within 30 min from the point of condensation. To minimise the confounding effects from interfaces, the confocal volume was placed at least 0.8 µm above and below from the coverslip-condensate interface and the top of condensate-water interface, respectively. Trace-labelled PGL-3 were used to observe the position and interfaces of condensate. Microscopy was performed as described above.

*Analysis:*

Image analysis was performed semi-automatically using custom scripts. First, an ROI was manually created to include only the inside of condensates but excluding the area at least 1 µm inwards from the contour of condensates in order to remove microspheres close to the condensate-water interface. Areas with microsphere aggregates were also excluded from ROIs. Automatic single particle tracking was performed using a custom script and the TrackMate plugin in Fiji [4,5]. Following the tracking, each movie was manually checked for inappropriate data as follows. Trajectories involving microsphere aggregates were further removed. Any movies with apparent linear movement due to condensate fusion or wetting events were either entirely discarded, or only the affected frames were disregarded for analysis. Using custom scripts, we calculated ensemble time-averaged mean squared displacement (MSD), meaning all trajectories from each condensate were pooled together without time dependence. Diffusion coefficient (D) and exponent α were calculated by fitting MSD to the 2D diffusion model $MSD = 4Dt^{\alpha}$. Using Stokes-Einstein-Sutherland equation $D = k_BT/6\pi\eta r$, we estimated the viscosity η. The reported values are mean ± 95% CI.

**Crosslinking mass spectrometry (XL-MS)**

*Sample preparation:*

Full-length PGL-3 was thawed and centrifuged at 21,000g for 5 min to remove large particles. The protein solution was diluted to 7.5 µM PGL-3, 125 mM (permissive for phase separation) or 500 mM NaCl (non-permissive for phase separation ), 25 mM HEPES pH 7.5 at a 100 µL scale. The diluted protein solution was incubated at room temperature for 5 min before adding 2 µL of disuccinimidyl dibutyric urea (DSBU, Thermo Fisher) from a 100 mM stock in DMSO (f.c. 2 mM). The reaction was incubated at room temperature on an end-over-end rotator for 1 h (10 rpm). Crosslinking reaction was quenched by addition of Tris-HCl pH 7.5 (f.c. 20 mM), and incubated at room temperature for 30 min.

Crosslinked samples were denatured by addition of solid urea (96.1 mg, f.c. 8 M, Sigma-Aldrich) and ammonium bicarbonate pH 8.0 (Ambic, f.c. 100 mM, Acros Organics), and diluted to a final volume of 200 µl with Optima LC/MS grade water. The samples were reduced by the addition of dithiothreitol (DTT, f.c. 5 mM, Sigma-Aldrich) and incubation for 30 min at 30 °C with agitation (700 rpm) on a benchtop thermomixer (Eppendorf). Samples were alkylated by the addition of iodoacetamide (IAA, f.c. 15 mM, Acros Organics) for 45 min at room temperature without agitation in the dark. DTT (f.c. 5 mM) was then added to quench excess IAA and the samples were incubated for 5 min at room temperature. The samples were diluted by the addition of 3 volumes of 100 mM Ambic pH 8.0 to reduce the final concentration of urea to 2 M. The samples were digested by trypsin (1:50 enzyme:protein w/w ratio, Pierce) and incubated at 25 °C, 700 rpm, overnight (~16 h) on a thermomixer. Next day, the digested samples were acidified with

Optima LC/MS grade 1% trifluoroacetic acid (TFA, Fisher Chemical) and desalted using C18 sep-pak cartridges (Waters). The acidified digests were diluted to 1 mL final volume using Buffer A [0.5% TFA in Optima LC/MS water]. C18 cartridges were placed on a vacuum manifold, conditioned twice using 1 ml Buffer B [0.5% TFA, 80% Optima LC/MS grade acetonitrile], followed by equilibration (four times with 1 mL) with Buffer A. Acidified digests were loaded onto the column under a reduced vacuum (1 mL/min), and then washed with Buffer A (four times with 1 mL ). The cartridges were placed on a 15 mL falcon tubes and samples were eluted by centrifugation at 350 rpm for 5 min in a 5910R centrifuge (Eppendorf). The samples were then transferred to a fresh tube and then reduced to dryness using a Vacufuge centrifugal concentrator (Eppendorf). Dried samples were stored at -80 °C until further analysis.

*LC-MS/MS:*
The LC-MS/MS experiments utilized an UltiMate3000 UHPLC system (Thermo Fisher) coupled with a Q-Exactive HF-X Orbitrap mass spectrometer (Thermo Fisher). To prepare samples, the dried peptides were vigorously resuspended in a 0.1% formic acid in Optima LC/MS grade water by vortexing and sonication. The peptide concentration was measured with Nanodrop One$^{C}$ microvolume UV–vis spectrophotometer (Thermo Fisher Scientific), and typically ~1 µg of the peptide was injected onto the LC-MS/MS system for each sample in triplicates. The remaining LC-MS/MS methods and parameters follow as described previously [6].

*Crosslink search*:
Scout [7] v. 1.5.1 was used to search for the crosslinked peptides using a database comprising just the PGL-3 full-length sequence. DSBU_KSTY crosslinker was selected in Scout with default settings. The peptide length was set to minimum 6 and maximum 60 residues. The minimum peptide mass was set to 500 Da and maximum peptide mass was set to 6000 Da. The precursor mass tolerance was set to 10.0 ppm, and the fragment ion precision was set to 20.0 ppm. Digestion enzyme was set to be fully specific trypsin, with cleavage sites after Lys and Arg and cleavage blocked by Pro, and a maximum of three missed cleavages were allowed. DSBU crosslinker mass of 196.0848 Da and MS-cleavable fragments of amine (light, 85.0527 Da) and isocyanate (heavy, 111.0320 Da) with mass shift of 25.9792 Da were set, with residue specificity towards KSTY residues. Static modifications included the carbamidomethylation of cysteine (Delta mass 57.0214 Da), and oxidation of methionine was used as variable modification (Delta mass 15.9949 Da). The 2% FDR cut-off was set to filter the crosslink spectral matches (CSMs) at the peptide and residue pair levels. All identifications are summarized in Dataset S1.

*Classification of crosslinks compatibility to published monomer/dimer structures:*
Because the previous works reported D1 and D2 dimer structures of *C. elegans* PGL-1, PGL-3 SWISS-MODEL models were used to map detected crosslinks to structures [8]. The SWISS-MODEL models were templated on 5w4a.1.A[9], for D1 dimer and 5cv1.1[10] for D2 dimer. PGL-1 and PGL-3 are highly conserved with 68.4% identify and 84.0 % similarity for D1, and 69.5 % identify and 84.4 % similarity for D2(Fig. S1A, B). The RMSD of published PGL-1 structures and PGL-3 SWISS-MODEL structures were very low, (0.066 for D1 and 0.069 for D2 (Fig. S4A)). In PyMOL using custom scripts, Cα-Cα distances for all crosslinks were measured on the monomers and dimers to obtain intra- and inter-chain distances, respectively. Then, crosslinks below or equal to 30 Å in distance were classified 'compatible' with the tested structure and the rest 'incompatible'. We used 30Å as a cut off based on the length of DSBU. This analysis yields five categories for crosslinks: 'compatible only with intra-chain (i.e., both crosslink sites on the same chain), 'compatible only with inter-chain' (i.e., crosslink sites on different chains),

’compatible with both intra and inter chains’ (i.e., crosslink sites on either the same or different chains), ’not compatible with either’, and ‘no structural information’.

*Crosslink abundance and frequency analysis:*
In Fig. 4A, all unique residue pairs are shown. In Fig. 4B, the total number of crosslinked peptide-spectrum matches (XSMs) associated with crosslinks between the indicated domains are shown, summing across the three replicates. In Fic. 4C, the number of XSMs associated with crosslinks between the indicated domains was divided by the total number of XSMs to generate frequencies. In Fig. S4B, the unique residue pairs are shown that were identified in both conditions or only in one.

**Mass photometry**
Mass photometry was performed on TwoMP (Refeyn) using AcquireMP and DiscoverMP (Refeyn) software. Unlabelled PGL-3(D1-D2) was centrifuged at 21,000 g for 5 min remove large particles and then diluted to four fold higher concentrations of final protein concentrations in 125 mM NaCl, 25 mM HEPES pH 7.5. The diluted protein was incubated at room temperature for at least 5 min. The mass photometry experiment was performed as described by manufacture protocol. Briefly, 12 µl of 125 mM NaCl, 25 mM HEPES pH 7.5 was placed on imaging well to focus and 4 µl of diluted proteins were mixed in by pipetting up and down. The video recording was started briefly after the final dilution. Mass calibration was performed similarly, using sweet potato β-amylase (sigma).

**A**

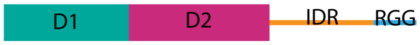


```
WATER
# Length: 748
# Identity:     444/748 (59.4%)
# Similarity:   543/748 (72.6%)
# Gaps:          75/748 (10.0%)

                              10        20        30        40        50        60        70        80        90
PGL-1             1 MEANKREIVDFGGLRSYFFPNLAHYITKNDEELFNNTSQANKLAAFVLGASKDAPGDEDILEMILPNDANAAVIAAGMDVCLLLGDKFRPKFDAAAEKLS    100
                    ||||||:||:..|::|||||:||||:..|||.|.||.:||||||||||||:...|.:|:|.||||||::|.|:||||||||:|||.||||||:.|||||
PGL-3             1 MEANKRQIVEVDGIKSYFFPHLAHYLASNDELLVNNIAQANKLAAFVLGATDKRPSNEEIAEMILPNDSSAYVLAAGMDVCLILGDDFRPKFDSGAEKLS    100
                              10        20        30        40        50        60        70        80        90

                              10        20        30        40        50        60        70        80        90
PGL-1           101 GLGHAHDLVSVIDDDKKLGMLARKAKLKKTEDAKILQALLKVIAIDDAAEKFVELTELVSQLDLDFDVYVLTKILGLISEETSDEVDIIRDNVVNAFDSC    200
                    .||.||||..:|||:||:.|||||.||||:.||||||.||||:..::|.||||||:||.|.||||||||||.|:||..|||..:|::||||||.:||::|
PGL-3           101 QLGQAHDLAPIIDDEKKISMLARKTKLKKSNDAKILQVLLKVLGAEEAEEKFVELSELSSALDLDFDVYVLAKLLGFASEELQEEIEIIRDNVTDAFEAC    200
                              10        20        30        40        50        60        70        80        90

                              10        20        30        40        50        60        70        80        90
PGL-1           201 KPLLKQLMLDGPKSEPADPFISLLMDPLEESVGKVVNHIAQLFEEASKNEGDESLVLRSQLGYQLFFLIVRSLADGKREVSKKILSGIPTSVRAEVFPGL    300
                    |||||:||::|||.:..|||..||:.|.|||:.|.|:||...|||||..|.||||||:|||||||.||:|||||||||:.|:.|.|.:|:||||||||||
PGL-3           201 KPLLKKLMIEGPKIDSVDPFTQLLLTPQEESIEKAVSHIVARFEEASAVEDDESLVLKSQLGYQLIFLVVRSLADGKRDASRTIQSLMPSSVRAEVFPGL    300
                              10        20        30        40        50        60        70        80        90

                              10        20        30        40        50        60        70        80        90
PGL-1           301 QRSVYKSAVFLGNHIIQVLLGSKKSFEDWDVVGVAKDLESAWKRRAIAELIKKFQVSILEQCFDKPVPLIPQSPLNNDAVIDNVNKALQFALWLTEFYGS    400
                    ||||:||||||.:|||||.|||.||||||..||:|:||||.|:|||||||:|||::|:|||||.:|:||:|||.|||:.||:|||.|||||||:||||||
PGL-3           301 QRSVFKSAVFLASHIIQVFLGSMKSFEDWAFVGLAEDLESTWRRRAIAELLKKFRISVLEQCFSQPIPLLPQSELNNETVIENVNNALQFALWITEFYGS    400
                              10        20        30        40        50        60        70        80        90

                              10        20        30        40        50        60        70        80        90
PGL-1           401 ENETEALGELRFLDSTSKNLLVDSFKKFVQGINSKTHVTRIVESLEKCCLSDTPSGRKSNVQPSTSQQQDSAYTKEEMTTVHNTYSVNTKAQVLNGLSDT    500
                    |:|.::|.:|:||...|||||||||||.||::||.||.||:|||||          .|:.:||.        |.::.||.:...:|:|.|||
PGL-3           401 ESEKKSLNQLQFLSPKSKNLLVDSFKKFAQGLDSKDHVNRIIESLEK----------SSSSEPSA--------TAKQTTTSNGPTTVSTAAQV-------    475
                              10        20        30        40        48                  60        70

                              10        20        30        40        50        60        70        80        90
PGL-1           501 NSSGLLVDSKDSLSLQEISCDEVDSSTLLSSSRNIGEGVTVKAVDPVPEKVNDAQQQQTVNEIEMASDANQDTSSSASPEVAPSFSTDGWDSPTKSVALP    600
                         :...|...|.|.|.|:..|.:.:|:|::.|||.|||.|.|.:|||:|              ::.|.:|.|:|||   ..||:||||||||||||
PGL-3           476 -----VTVEKMPFSRQTIPCEGTDLANVLNSAKIIGESVTVAAHDVIPEKLN--------------AEKNDNTPSTASP---VQFSSDGWDSPTKSVALP    553
                              80        90        00        10        20                       30        40        50

                              10             20        30        40       50         60        70        80        90
PGL-1           601 PGMQQIDEEETTVADKDST-----PQPQARAETAWGSGDATPMPLPAPTNQYKVSGFGEAKVAKGFGQFAPTSSAYGGGGGRGGYG---GGDRGGRGGYG    692
                    |.:..::||:    ::|:|     ||||.|..|||||||||:||..|.|:|||||||.|.||.||||||.::   |..|||.||   ||||||||.||
PGL-3           554 PKISTLEEEQ----EEDTTITKVSPQPQERTGTAWGSGDATPVPLATPVNEYKVSGFGAAPVASGFGQFASSN----GTSGRGSYGGGRGGDRGGRGAYG    645
                           60                  70        80        90        00        10        20             30        40

                                  00        10             20
PGL-1           693 GDRG------GRGGYGGGDRGGRGGYG-GDR----GRGGYGGRGGRGGF    730
                    ||||      |..||.||||||||.|| |.|    ||.|:.| |.|||
PGL-3           646 GDRGRGGSGDGSRGYRGGDRGGRGSYGEGSRGYQGGRAGFFG-GSRGGS    693
                        50        60        70        80        90
```

**B**

| | Identity | Similarity | Gap |
|---|---|---|---|
| FL | 444/748 (59.4%) | 543/748 (72.6%) | 75/748 (10.0%) |
| D1 | 145/212 (68.4%) | 178/212 (84.0%) | 0/212 ( 0.0%) |
| D2 | 169/243 (69.5%) | 205/243 (84.4%) | 0/243 ( 0.0%) |
| IDR | 90/208 (43.3%) | 122/208 (58.7%) | 38/208 (18.3%) |
| RGG | 43/71 (60.6%) | 44/71 (62.0%) | 15/71 (21.1%) |
| D1D2 | 309/447 (69.1%) | 375/447 (83.9%) | 0/447 ( 0.0%) |
| IDR-RGG | 133/283 (47.0%) | 166/283 (58.7%) | 57/283 (20.1%) |

**Fig. S1.** (**A**) Alignment of PGL-1 and PGL-3, using EMBOSS Water [11]. (**B**) Table showing similarity and identity of protein domains between PGL-1 and PGL-3.

**C**

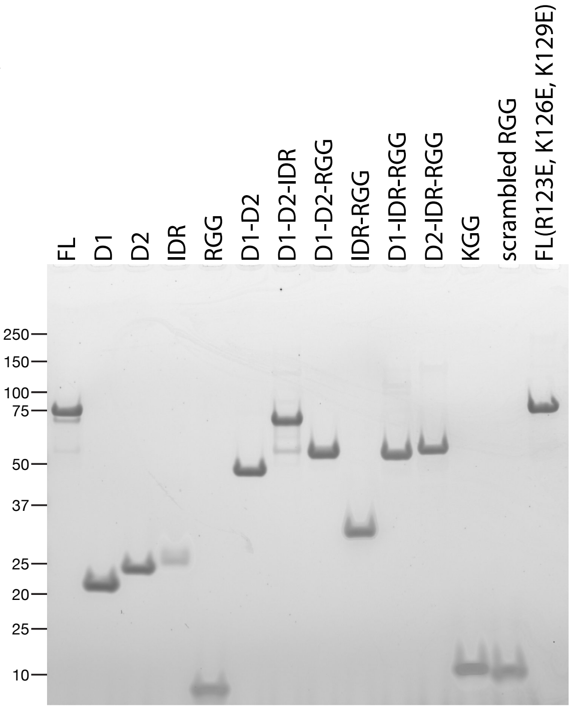


**D**

| Construct | Residues | MW (kDa) |
| --- | --- | --- |
| FL | 1-693 | 74.8 |
| D1 | 1-212 | 23.5 |
| D2 | 205-452 | 28.0 |
| IDR | 448-622 | 18.1 |
| RGG | 623-693 | 6.76 |
| D1-D2 | 1-447 | 50.0 |
| D1-D2-IDR | 1-622 | 68.1 |
| D1-D2-RGG | 1-447::623-693 | 56.78 |
| IDR-RGG | 448-693 | 24.8 |
| D1-IDR-RGG | 1-212::448-693 | 48.3 |
| D2-IDR-RGG | 205-693 | 51.9 |
| KGG | 623-693(all R to K) | 6.39 |
| scrambled RGG | 623-693(scrambled) | 6.76 |

**E**

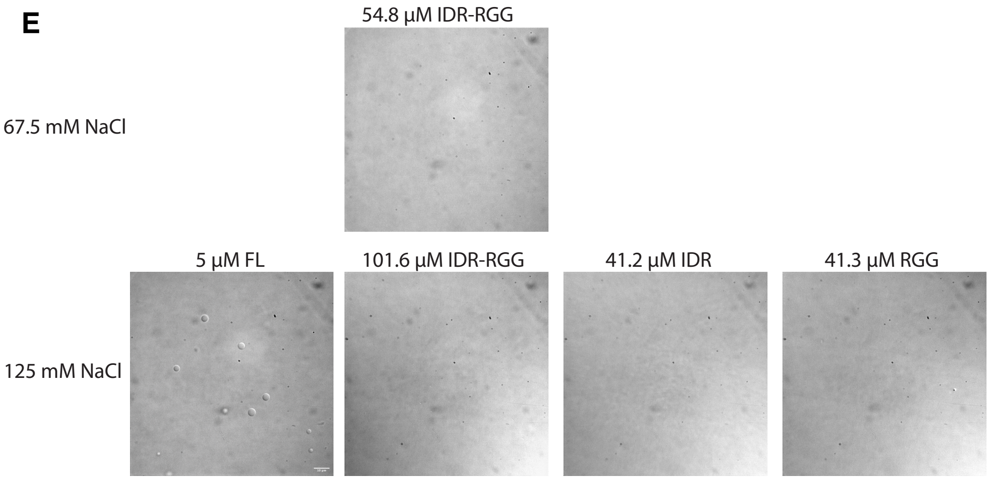


**Fig. S1.** (**C**) Protein gel stained with Coomassie G-250 showing the recombinant PGL-3 derivatives used in the condensation assays shown in Fig. 1C, D. 1 µg protein per well was loaded. **(D)** Table listing the region and the molecular weight for each PGL-3 constructs used in this study. (**E**) DIC micrographs of unlabelled PGL-3 and its derivatives of indicated protein concentrations at 67.5 or 125 mM NaCl. Scale bar=10 µm.

**F**

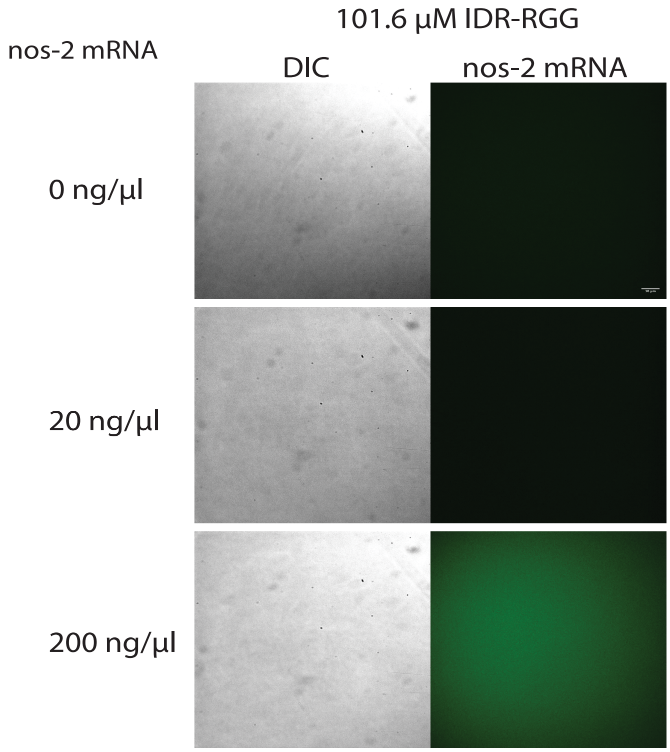


**G**

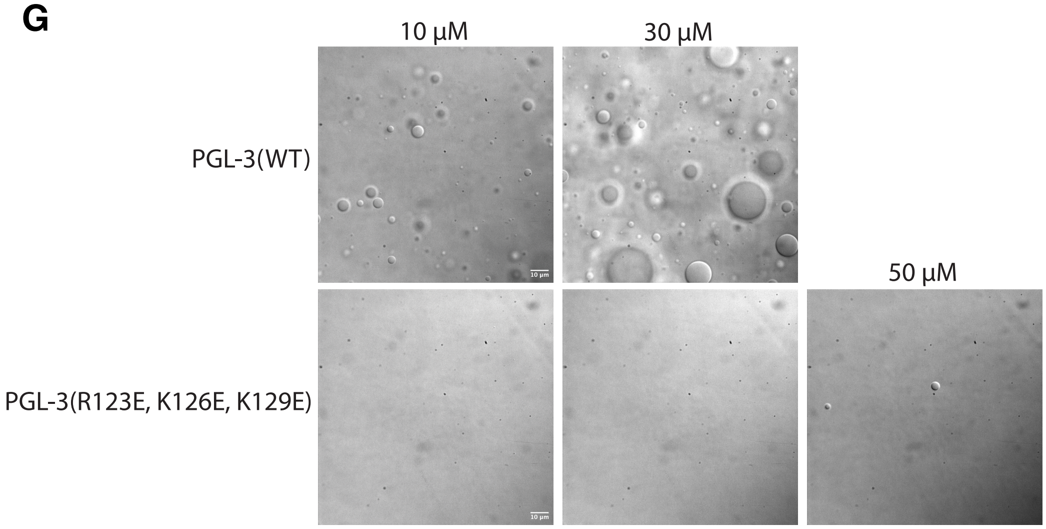


**Fig. S1. (F)** DIC micrographs of unlabelled PGL-3 IDR-RGG at 101.6 μM PGL-3(IDR-RGG), 125 mM NaCl and varying concentrations of *nos-2* mRNA. Scale bar=10 μm. **(G)** DIC micrographs of unlabelled wild-type and D1 dimerization mutant of full-length PGL-3 tested for condensation at the indicated protein and salt concentrations. Note the robust condensation of wild-type PGL-3, which is not observed for the mutant. PGL-3(R123E, K126E, K129E) is full-length PGL-3 with mutations in the D1 domain, previously shown to interfere with dimerization in PGL-1 [9]. Scale bar=10 μm.

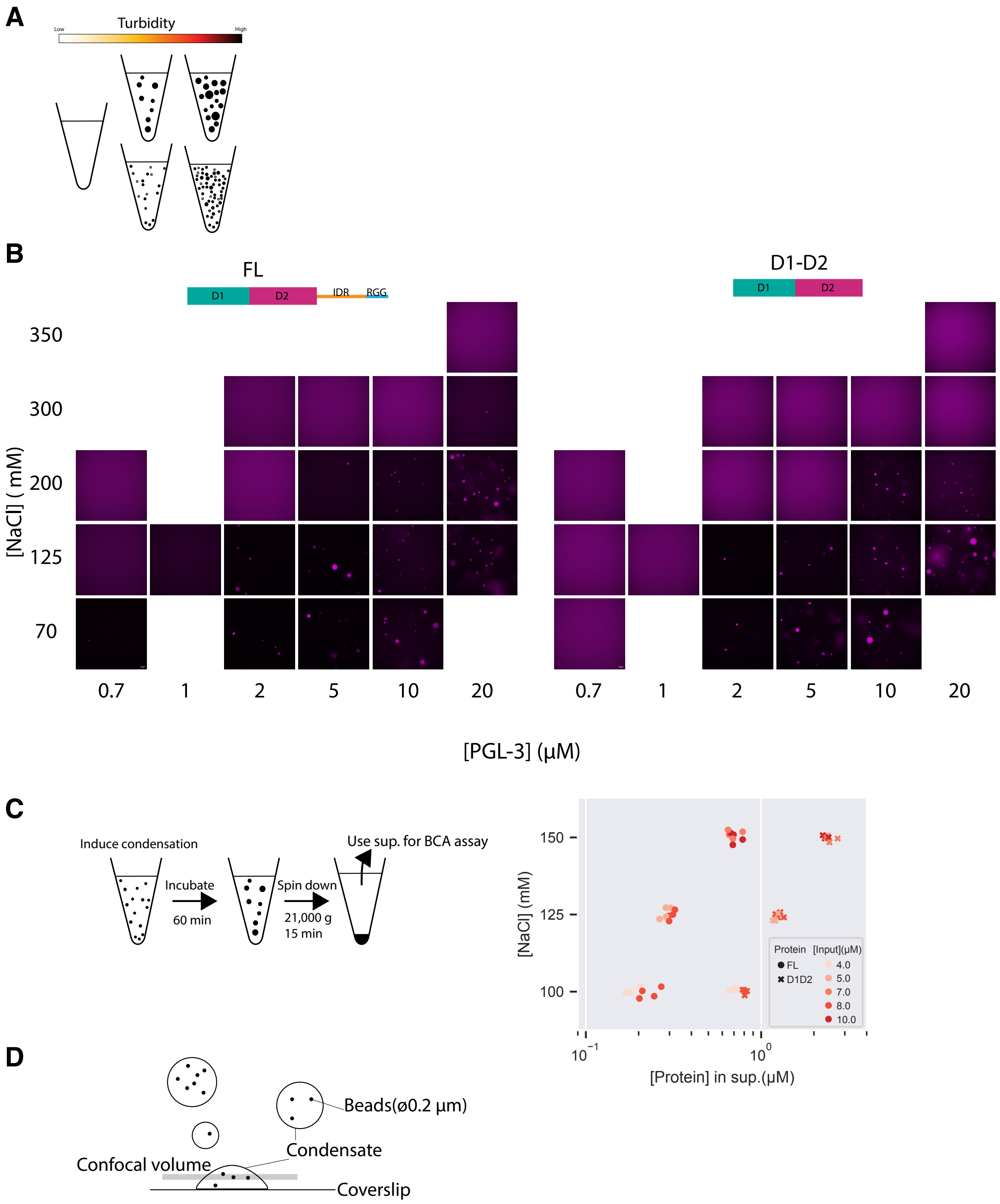


**Fig. S2.** (**A**) Schematic describing the basis of turbidity measurement. Related to Fig. 2A. (**B**) Representative fluorescence micrographs at indicated protein and salt concentrations for full-length PGL-3 and D1-D2. Scale bar=10 μm. (**C**) Left: Schematic describing the procedure of $c_{sat}$ estimation by $c_{dil}$ measurement. Right: Plot showing $c_{dil}$ of FL and D1-D2 at varying protein and salt concentrations, permissive for condensation. (**D**) Schematic describing the viscosity measurement experiments. Related to Fig. 2C.

| | $C_{dil}$ (mean± 95% confidence interval) |
|---|---|
| **FL** | 0.27 ± 0.03 |
| **D1-D2** | 0.94 ± 0.03 |
| **D1-D2-RGG** | 0.51 ± 0.02 |
| **D1-D2-3x(GGGGS)-RGG** | 0.48 ± 0.01 |

**Fig. S3.** Table showing $c_{dil}$ of PGL-3 constructs, related to Fig. 3B. Values show mean±95% confidence interval

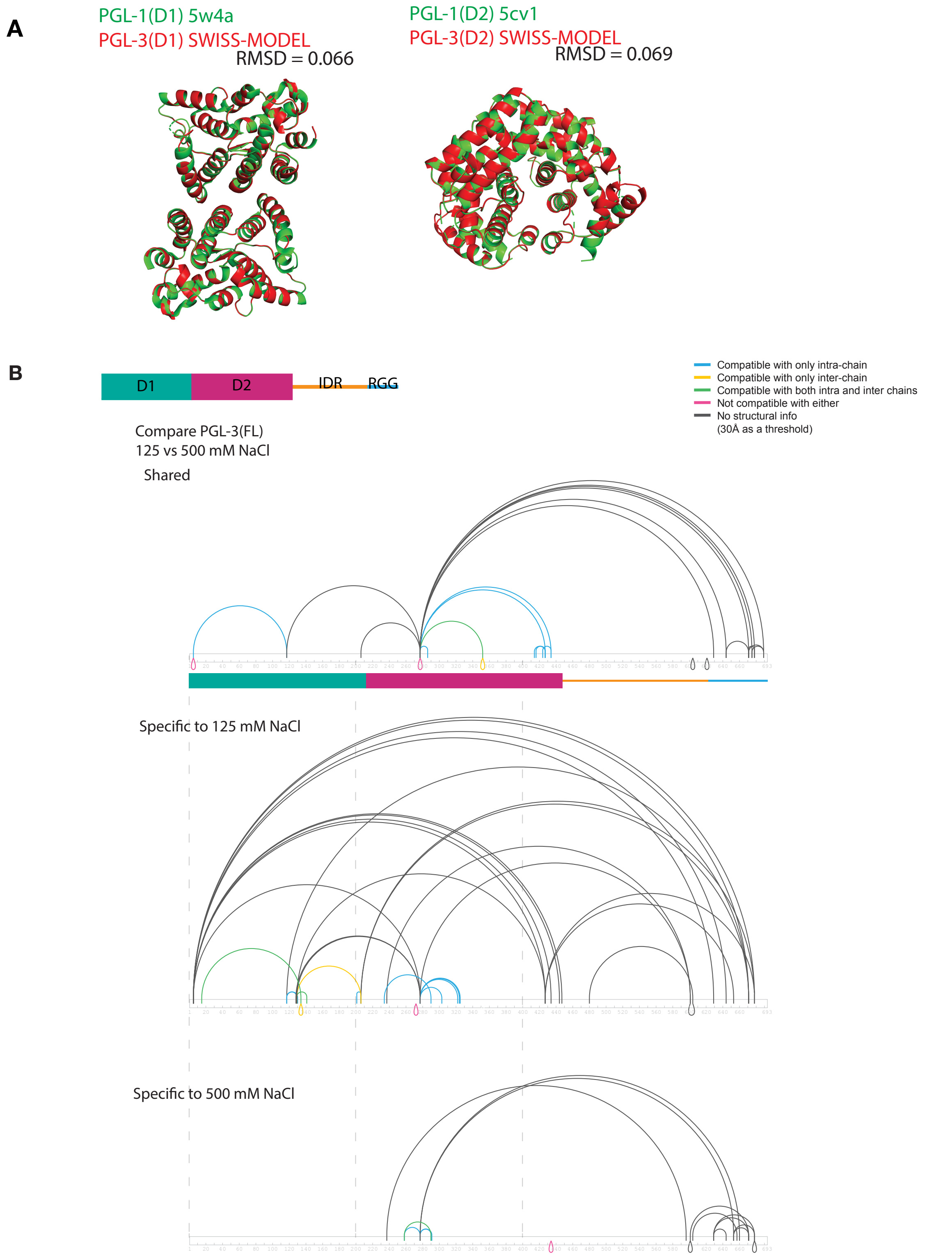


**Fig. S4.** (**A**) Alignment of D1(1-212 a.a.) and D2 (205-447 a.a.) between published crystal D1 and D2 dimer structures of PGL-1[9,10] and SWISS-MODEL[8] structures of PGL-3.
(**B**) Connectogram comparing crosslinking profiles of PGL-3 at 125 mM and 500 mM NaCl. Top: Shared between two conditions. Middle: Specific to 125 mM NaCl. Bottom: Specific to 500 mM NaCl.

**A**

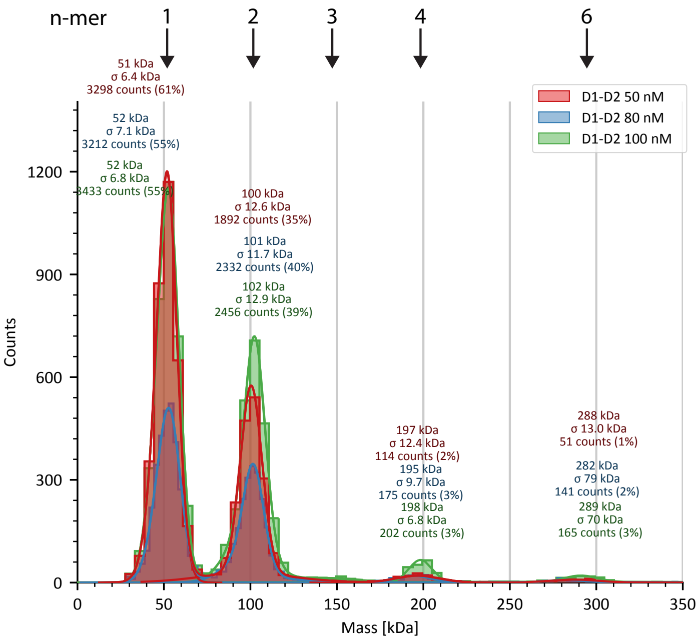


**B**

```
RGG      1  GTSGRGSYGGGRGGDRGGRGAYGGDRGRGGSGDGSRGYRGGDRGGRGSYG  50
KGG      1  GTSGKGSYGGGKGGDKGGKGAYGGDKGKGGSGDGSKGYKGGDKGGKGSYG  50
scrRGG   1  GGEGYFGRYFGYRGRSGRSGGSGGQGRGSGGSRDYGRGAGGRGRGSGGRD  50

RGG      51 EGSRGYQGGRAGFFGGSRGGS 71
KGG      51 EGSKGYQGGKAGFFGGSKGGS 71
scrRGG   51 SGRSGGGAGGGDTGGRGRDYG 71
```

**C**

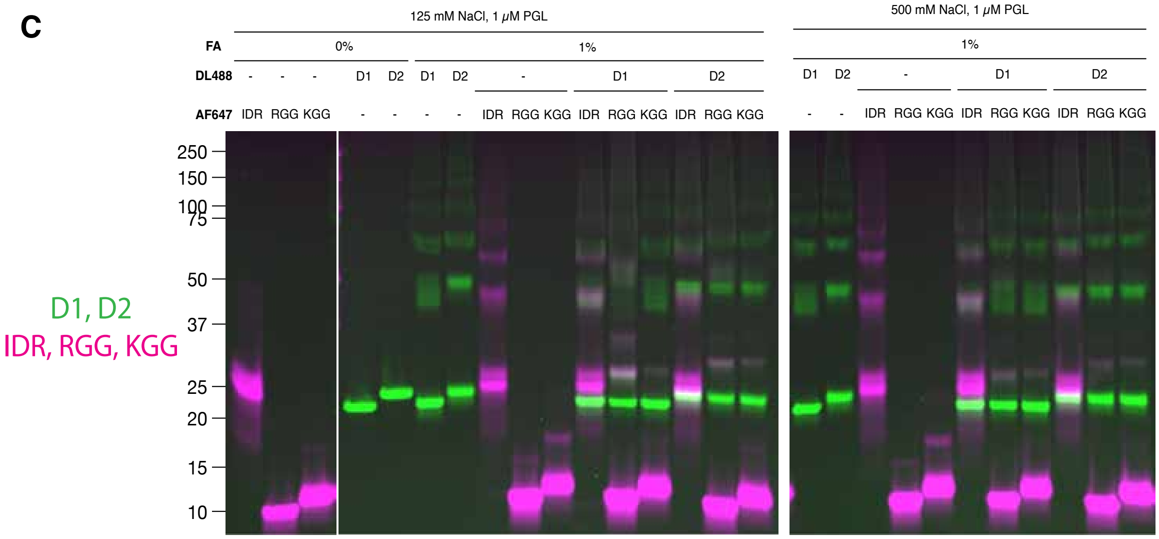


**Fig. S5.** (**A**) Graph showing the mass distribution of D1-D2 oligomers in 125 mM NaCl solution, measured by mass photometry. D1-D2 was diluted by 4-fold to reach the indicated concentrations in imaging wells and the videos were immediately recorded by TwoMP (Refeyn). (**B**) Alignment among the native RGG peptide, the KGG peptide and the scrambled RGG peptide. RGG triplets and RG doublets are highlighted in magenta and yellow, respectively. (**C**) Dual-color fluorescent image of the gel same as Fig. 5A. Two channels overlaid in color (DyLight 488 in green and Alexa Fluor 647 in magenta).

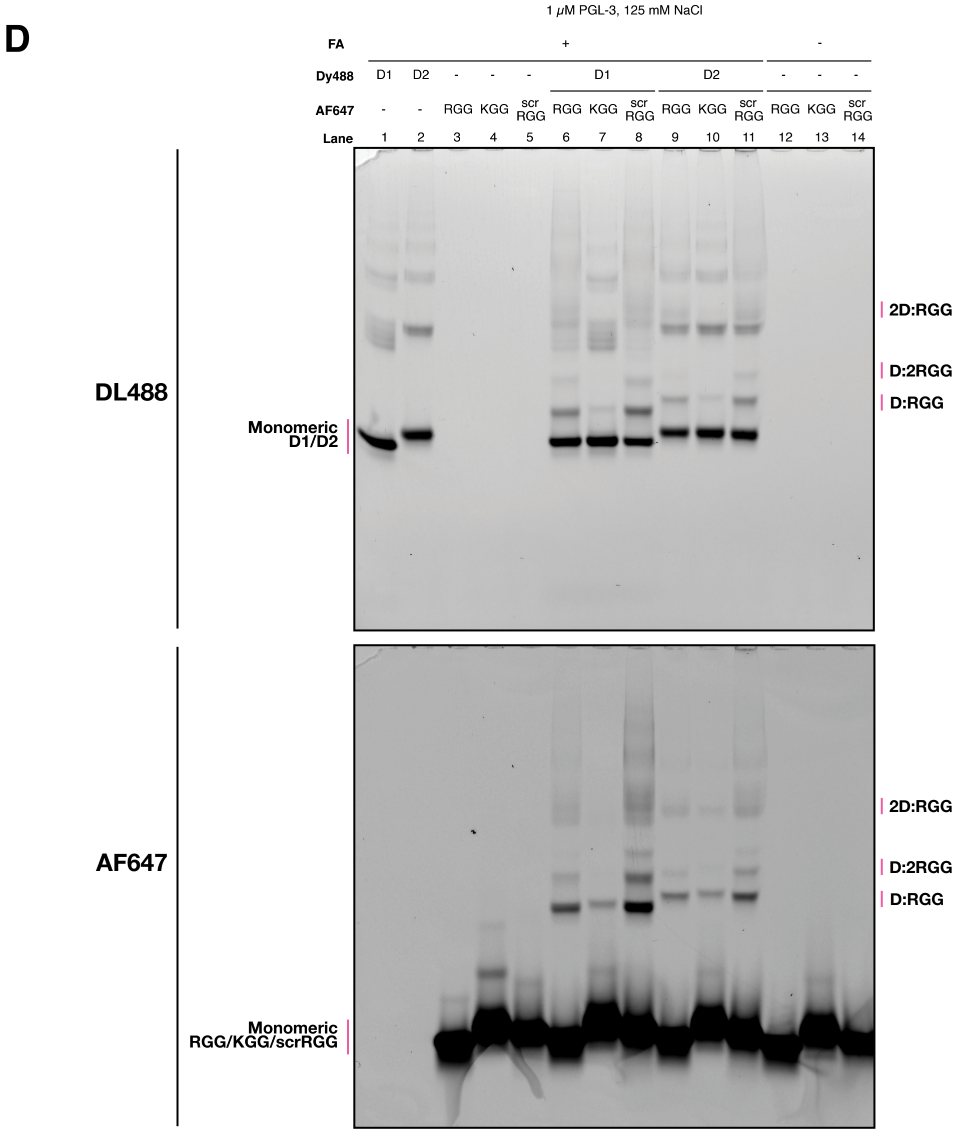


**Fig. S5.** (**D**) Fluorescent images of SDS-PAGE gels showing crosslinked complexes of PGL-3 D1 or D2 with RGG, KGG or scrambled RGG (scrRGG) peptides.

**E**

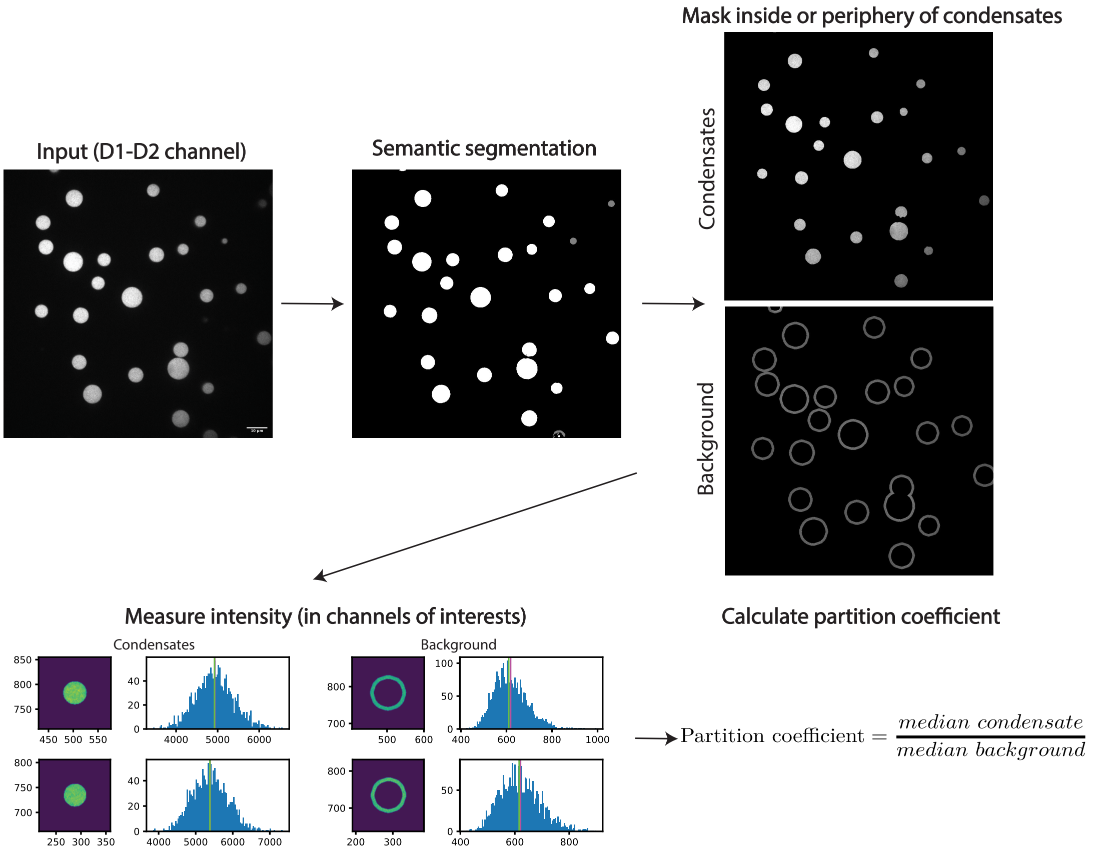


**F**

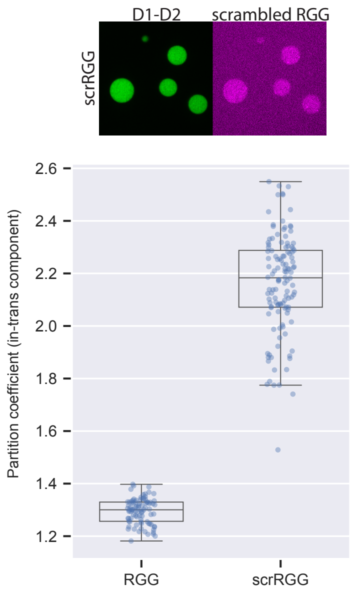


**G**

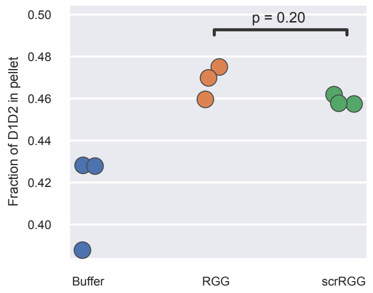


**Fig. S5.** (**E**) Schematics describing the algorithm for determining partition coefficient. Related to Figure 5B, C. (**F**) Top: Photomicrograph showing partition of scrambled RGG (magenta) to D1-D2 condensates (green). Bottom: Graph showing the partition coefficient of scrambled RGG to D1-D2 condensates. (**G**) Graph showing results from pelleting assay using scrambled RGG, where fractions of D1-D2 protein in pellet over total protein were calculated upon condensation of D1-D2 with the native or scrambled RGG and two phases were separated by centrifugation. P value by Wilcoxon rank-sum test.

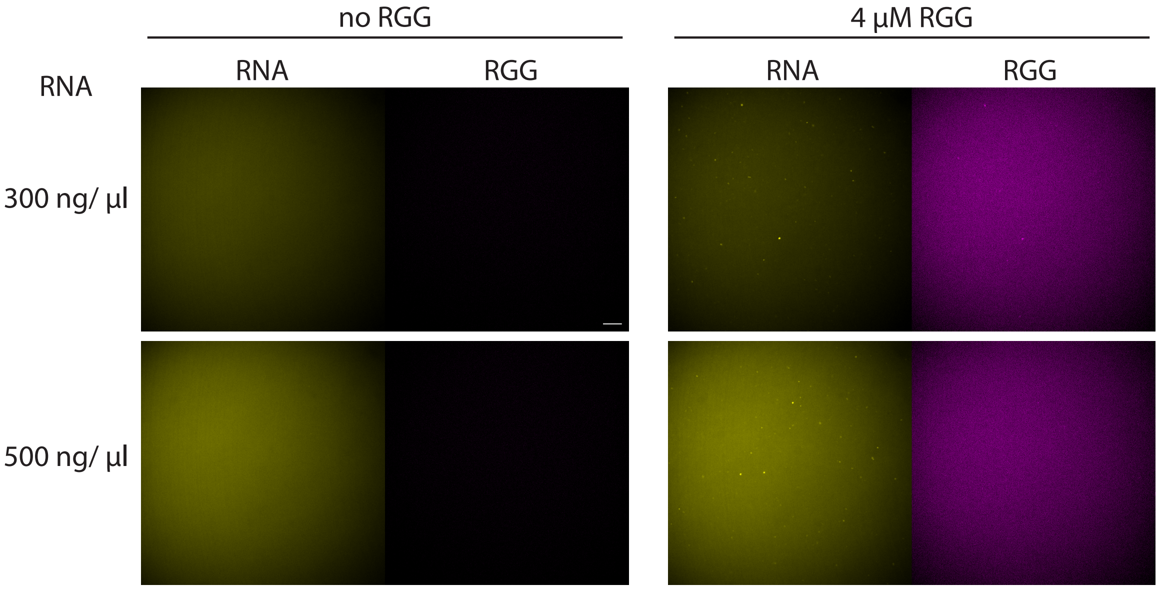


**Fig. S6.** Fluorescence micrographs showing distribution of *nos-2* mRNA (yellow) and the RGG peptide (magenta) at indicated concentrations, without D1-D2. Note RNA and RGG form co-aggregates only when the two components are mixed.

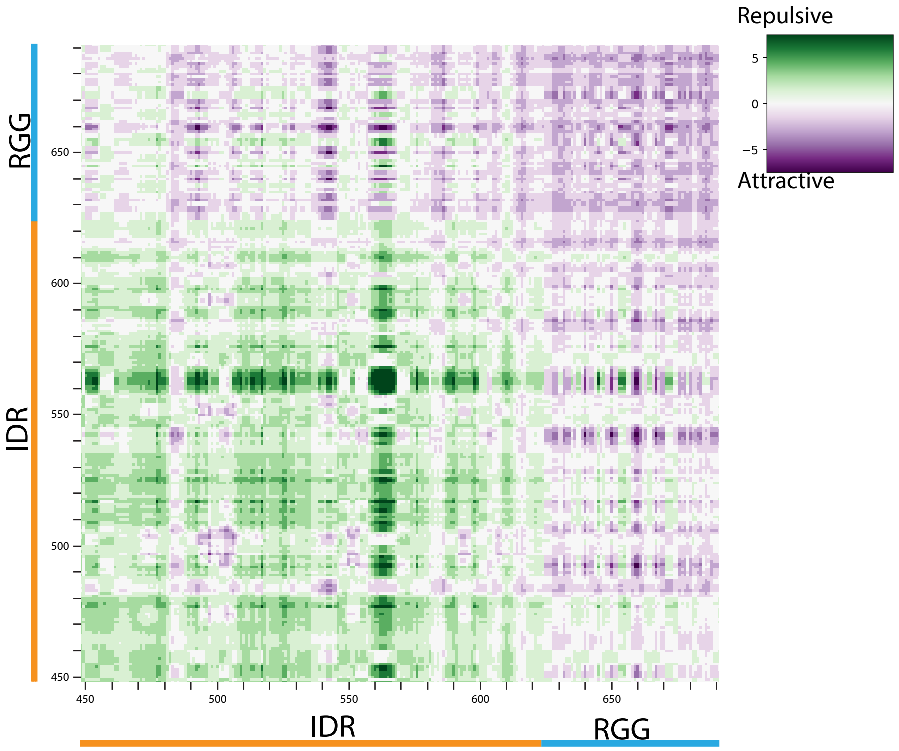


**Fig. S7.** FINCHES [12] intermap for IDR-RGG region of PGL-3. Green indicates repulsive and purple indicates attractive pairs.

**Table S1**. List of plasmids used to express PGL-3 and variants. All plasmids were prepared using the pMAL-c2X vector backbone.

| Plasmid | Construct | Tag (Removed by TEV cleavage during purification) | N-terminal linker (Remaining after TEV cleavage) | PGL-3 amino acids |
|---|---|---|---|---|
| pRK015 | PGL-3(FL) | 6xHis-MBP-6xHis-TEV | none | 1-693 |
| pRK049 | PGL-3(D1) | 6xHis-MBP-6xHis-TEV | none | 1-212 |
| pRK050 | PGL-3(D2) | 6xHis-MBP-6xHis-TEV | G | 205-447 |
| pRK011 | PGL-3(IDR) | MBP-6xHis-TEV | none | 448-622 |
| pRK036 | PGL-3(D1-D2) | 6xHis-MBP-6xHis-TEV | none | 1-447 |
| pRK042 | PGL-3(D1-D2-IDR) | 6xHis-MBP-6xHis-TEV | none | 1-622 |
| pRK082 | PGL-3(D1-D2-RGG) | 6xHis-MBP-6xHis-TEV | none | 1-447::623-693 |
| pRK084 | PGL-3(D1-D2-3(GGGGS)-RGG) | 6xHis-MBP-6xHis-TEV | none | 1-447::3x(GGGGS)::623-693 |
| pAAP35 | PGL-3(IDR-RGG) | MBP-6xHis-TEV | GAGL | 448-693 |
| pAAP36 | PGL-3(D1-IDR-RGG) | MBP-6xHis-TEV | GAGL | 1-212::448-693 |
| pAAP37 | PGL-3(D2-IDR-RGG) | MBP-6xHis-TEV | GAGL | 205-693 |
| pRK047 | PGL-3(FL)(R123E, K126E, K129E) | 6xHis-MBP-6xHis-TEV | none | 1-693(R123E, K126E, K129E) |

**Movie S1 (separate file).** Video showing trajectories of diffusing beads inside condensates. Left: full-length PGL-3, Right: D1-D2. Each video shows a close-up of beads inside one condensate. Scale bar = 5 μm. Related Fig. 2C.

**Dataset S1 (separate file).** Excel file showing the unique crosslinked residue pairs recovered in XL-MS under 125mM NaCl (second tab) and 500mM NaCl (third tab). The first tab describes the values for each column.
